# Evolutionary trade-offs in osmotic and ionic regulation and the expression of gill ion transporter genes in high latitude, Neotropical cold clime crabs from the ‘end of the world’

**DOI:** 10.1101/2022.02.11.480083

**Authors:** John Campbell McNamara, Anieli Cristina Maraschi, Federico Tapella, Maria Carolina Romero

**Author notes:** **Corresponding author:** Professor John Campbell McNamara, Departamento de Biologia, Faculdade de Filosofia, Ciências e Letras de Ribeirão Preto, Universidade de São Paulo, Ribeirão Preto 14040-901, SP, Brazil.

## Abstract

Seeking to identify consequences of evolution at low temperature, we examine hyper/hypo-osmotic and ionic regulation and gill ion transporter gene expression in two sub-Antarctic Eubrachyura from the Beagle Channel, Tierra del Fuego. Despite sharing the same osmotic niche, *Acanthocyclus albatrossis* tolerates a wider salinity range (2-65 ‰S) than *Halicarcinus planatus* (5-60 ‰S); respective lower and upper critical salinities are 4 and 12 ‰S, and 63 and 50 ‰S. *Acanthocyclus albatrossis* is a weak hyperosmotic regulator, while *H. planatus* hyper-osmoconforms; isosmotic points are 1,380 and ≈1,340 mOsm kg^−1^ H_2_O. Both crabs hyper/hypo-regulate [Cl^−^] well with iso-chloride points at 452 and 316 mmol L^−1^ Cl^−^, respectively. [Na^+^] is hyper-regulated at all salinities. mRNA expression of gill Na^+^/K^+^-ATPase is salinity-sensitive in *A. albatrossis*, increasing ≈1.9-fold at 5 ‰S compared to 30 ‰S, decreasing at 40 to 60 ‰S. Expression in *H. planatus* is very low salinity-sensitive, increasing ≈4.7-fold over 30 ‰S, but decreasing at 50 ‰S. V(H^+^)-ATPase expression decreases in *A. albatrossis* at low and high salinities as in *H. planatus*. Na^+^-K^+^-2Cl^−^ symporter expression in *A. albatrossis* increases 2.6-fold at 5 ‰S, but decreases at 60 ‰S compared to 30 ‰S. Chloride uptake may be mediated by increased Na^+^-K^+^-2Cl^−^ expression but Cl^−^ secretion is independent of symporter expression. These unrelated eubrachyurans exhibit similar systemic osmoregulatory characteristics and are better adapted to dilute media; however, the gene expressions underlying ion uptake and secretion show marked interspecific divergences. Cold clime crabs may have limited energy expenditure by regulating hemolymph [Cl^−^] alone, apportioning resources for other metabolic processes.

**Summary statement:** Sub-Antarctic crabs may skimp on osmoregulatory capabilities to apportion energy for metabolic processes. They regulate chloride but not sodium or osmolality. Transporter gene expressions diverge markedly. Adaptive, differential ion regulation may characterize cold clime crabs.

## Introduction

Remarkable physiological adaptations are often encountered in crustaceans that occupy extreme environments. During evolutionary diversification, such adaptations may have been driven by severe ambient selection pressures such as low temperatures and salinities that frequently have acted synergistically (Faria et al. 2017, 2020; Capparelli et al. 2021). The evolution of osmoregulatory capability is one such ancient physiological process that has enabled crustaceans to radiate from their ancestral marine habitat and to confront the challenges, not only of variable salinity environments, be they dilute or concentrated, such as encountered in intertidal, estuarine and semi-terrestrial habitats, but also of fresh water (McNamara and Freire, 2022). Within the brachyuran decapods in particular, many diverse patterns and degrees of osmoregulatory ability have been characterized (Mantel and Farmer 1983; Péqueux 1995). These include mechanisms of isosmotic intracellular regulation and cell volume adjustment on which osmoconforming marine crabs for example are wholly dependent (Freire et al. 2008b; Foster et al. 2010). They extend also to excellent freshwater hyper-osmoregulating crabs (Onken and McNamara 2002; Weihrauch et al. 2004; Mantovani and McNamara 2021) and aeglid squat lobsters (Faria et al., 2010; Freire et al., 2013; Bozza et al., 2019) and semi-terrestrial hyper-/hypo-osmoregulators (Leone et al. 2020) that effect anisosmotic regulation of their extracellular fluids using gill-based mechanisms of ion transport (McNamara and Faria 2012; Faria et al. 2017).

Many studies of decapod crustaceans have examined the molecular (Luquet et al. 2005; Faleiros et al. 2010, 2017; Havird 2014; Maraschi et al. 2021), biochemical (D’Orazio and Holliday 1985; Leone et al. 2015) and physiological mechanisms (Taylor and Taylor 1992; Péqueux 1995; Henry et al. 2012), and the ultrastructural rearrangements in the gill epithelia and antennal glands (McNamara and Torres 1999; Freire et al. 2008a; McNamara et al. 2015) that underlie the transport processes responsible for sustaining such osmotic and ionic gradients between the hemolymph and the external medium. However, most investigations have employed large, easy obtainable crabs and shrimps from northern hemisphere localities, particularly the Nearctic and Palearctic biogeographical zones, ranging from the shores of the tropical eastern Pacific and Atlantic Oceans through the coasts of temperate northern Pacific and Atlantic Oceans to Arctic climatic regimes (e. g., *Hemigrapsus oregonensis* and *H. nudus*, Dehnel 1962; *Rhithropanopeus harrisi*, Smith 1967; *Cancer irroratus* and *C. borealis*, Charmantier and Charmantier-Daures, 1991; *Carcinus maenas*, Siebers et al. 1982, Cieluch et al. 2004). In contrast, there are far fewer osmoregulatory studies on decapod Brachyura from the Neotropical (*Minuca rapax*, Zanders and Rojas 1996; *Neohelice granulata*, Genovese et al., 2004, Bianchini et al. 2008; *Callinectes danae*, Leone et al. 2015; *Uca, Minuca* and *Leptuca*, Thurman et al. 2017, Faria et al. 2017; *Ucides cordatus*, Leone et al. 2020), Afrotropical, and Australasian biogeographical regions (e. g., *Helice crassa*, Bedford, 1972; *Uca formosensis, U. arcuata, U. vocans* and *U. lactea*, Lin et al. 2002; *Hemigrapsus crenulatus* and *H. sexdentatus*, Falconer et al. 2019). Fewer yet have examined species from southern Patagonian shores, particularly the Magellanic zoogeographical province, including Tierra del Fuego at the southernmost tip of South America.

We have investigated critical thermal limits (Faria et al. 2017) and aerobic and anaerobic metabolism (Faria et al. 2020) in several families of Neotropical Eubrachyura distributed latitudinally from the Equator to sub-Antarctic latitudes. While the systemic oxygen consumption and enzyme kinetic responses of the tropical and subtropical crab species are similar despite their phylogenetic diversity, they differ markedly from those of the sub-Antarctic Magellanic crabs that show lower aerobic metabolic demands and higher rates of hemolymph lactate formation (Faria et al. 2020). Given the temperature-dependent energy demands of active ion transport, the evolution of physiological osmoregulatory processes in crabs from the sub-Antarctic zone may have incorporated a limiting, cold clime effect compared to the regulatory abilities seen in Brachyura from tropical and subtropical climes in the Southern Hemisphere. The hypothesis we evaluate here is that such limitations might be manifest in quantitative and/or qualitative alterations in osmotic and ionic regulatory abilities at different levels of structural organization in sub-Antarctic crabs.

However, there have been no osmoregulatory studies on sub-Antarctic crabs, and the very limited data available concern Mg^2+^ regulation alone in just two species, the belliid *Acanthocyclus albatrossis*, and the hymenosomatid *Halicarcinus planatus*. Hemolymph Mg^2+^ titers in *A. albatrossis* (21.6 mmol l^−1^) and *H. planatus* (10.7 mmol l^−1^) are hypo-regulated at very low concentrations compared to most Brachyura (30-50 mmol l^−1^) (Frederich et al. 2001). In contrast, the biogeography (Diez et al. 2011), distribution and ecology (López-Farrán et al. 2021), population dynamics and growth (Diez et al. 2013) and thermal and reproductive physiology (Diez et al. 2010) of these two sympatric species are well known, and reveal that the abundant populations of both species encountered in the Beagle Channel, Tierra del Fuego, occupy very similar osmotic and thermal niches.

Given the complete lack of osmoregulatory findings on crab species from high Neotropical, sub-Antarctic latitudes, we have conducted a detailed study of hyper- and hypo-osmotic and ionic regulation and gill ion transporter gene expression in these two common, sympatric sub-Antarctic crabs *A. albatrossis* and *H. planatus*, seeking to identify putative, energy-saving osmoregulatory traits. We reveal clear differences in their salinity tolerances, in hemolymph osmotic, Na^+^ and Cl^−^ regulatory abilities, and in their gill Na^+^/K^+^-ATPase and V(H^+^)-ATPase and Na^+^-K^+^-2Cl^−^ symporter mRNA expression on rigorous salinity challenge. Although phylogenetically distant, *A. albatrossis* and *H. planatus* exhibit similar systemic osmoregulatory characteristics and are better adapted to dilute than concentrated media, likely a result of convergent adaptation owing to the effect of low temperature on osmoregulatory capability. However, the gene expressions underlying ion uptake and secretion show marked interspecific divergence. Apparently, these crabs have limited their osmoregulatory energy expenditure by regulating hemolymph [Cl^−^] alone, apportioning energetic resources for other metabolic processes.

## Materials and Methods

### The study area

This study was conducted on the northern shores of the eastern Beagle Channel, Tierra del Fuego, Argentina, during the early southern autumn, from April to May of 2017. The collecting sites were located in a narrow stretch of the Channel, around 60 km east of Ushuaia and about 6 km north of Isla Navarino (Fig. 1).

**Figure 1.**
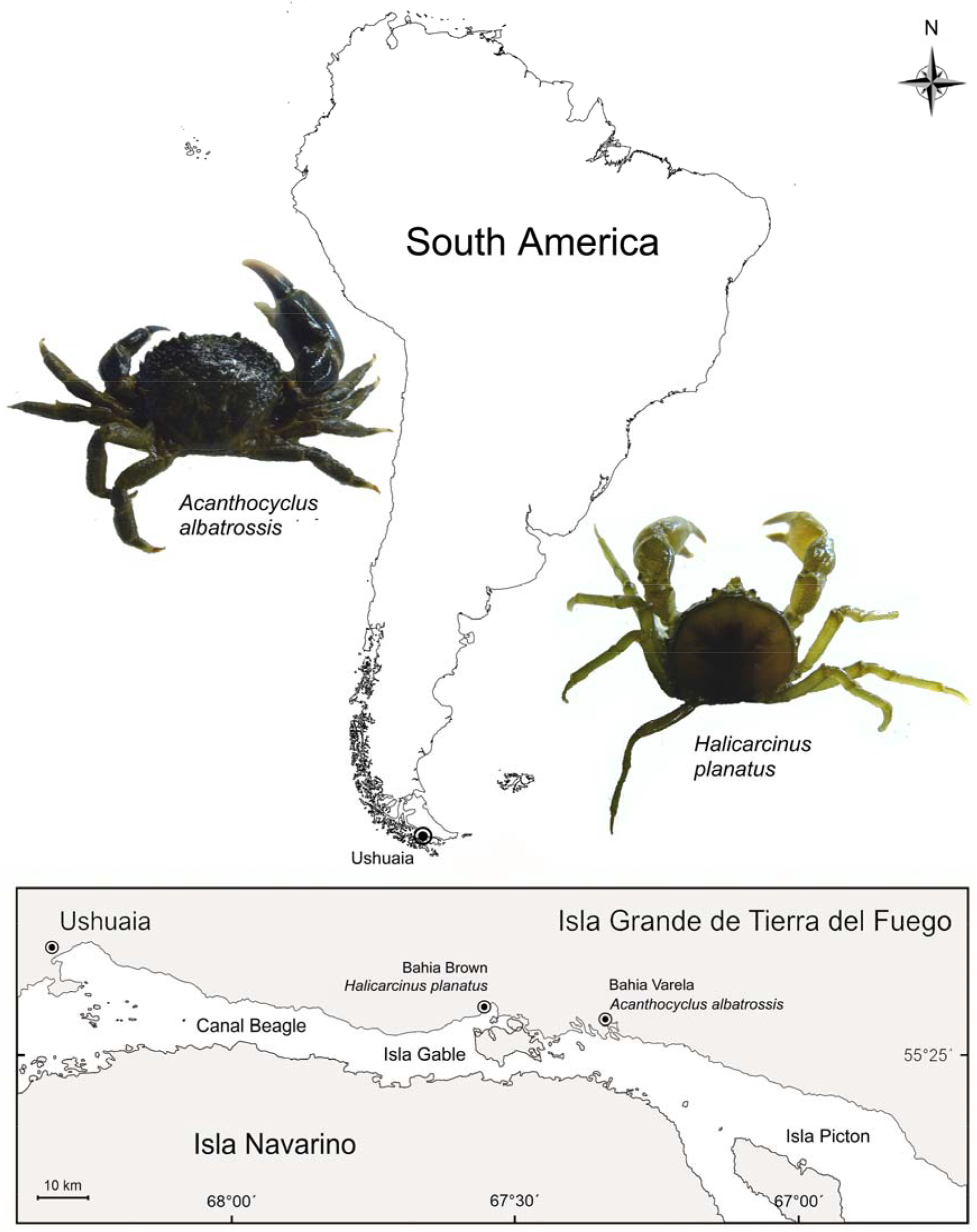
Location maps of South America and of the collecting sites for the Neotropical sub-Antarctic crabs *Acanthocyclus albatrossis* (Belliidae) and *Halicarcinus planatus* (Hymenosomatidae) in the Beagle Channel near Ushuaia, Tierra del Fuego, Argentina. Crabs were collected by hand at low tide from the infralittoral zone of stony beaches bordering Bahía Varela (54° 52’ 12.72” S, 67° 22’ 22.30” W) and Bahía Brown (54° 51’ 38.27” S, 67° 31’ 1.83” W), respectively, on the northern coast of the Beagle Channel.

Surface salinities in the mid Beagle Channel range from 29.4 to 32.7 ‰S year round (Isla et al. 1999; Diaz et al. 2018) while in coastal bays, salinity can be as low as 15.0 to 19.4 ‰S, owing to local fresh water runoff from rivers and streams, but reaching 31.7 ‰S due to evaporation, with an annual mean of ≈27 ‰S (Curelovich et al. 2009). The Beagle Channel thus exhibits salinities slightly less than seawater, and can be considered an estuarine regime in character (Isla et al. 1999).

Sea surface temperatures in the lower intertidal zone of the Beagle channel shores range from 4.2-5.2 °C during the southern winter (August) to 9.5-9.8 °C in summer (January) (Diez and Lovrich, 2010; Diez et al. 2013).

### Crab collections and laboratory maintenance

Approximately 300 specimens each, either male or female, of the crabs *Acanthocyclus albatrossis* (Belliidae) and *Halicarcinus planatus* (Hymenosomatidae) were collected by hand at low spring tide from beneath pebbles in the infralittoral zone (≈0.5 °C, ≈30 ‰S) of stony beaches on the shores of Bahía Varela (54° 52’ 12.72” S, 67° 22’ 22.30” W) and Bahia Almirante Brown (54° 51’ 38.27” S, 67° 31’ 1.83” W) located in the Beagle Channel, Tierra del Fuego, Argentina (Fig. 1).

Crab carapace widths were ≈3 cm and ≈2 cm for *A. albatrossis* and *H. planatus*, respectively. The crabs were transported by utility vehicle (1.5 h, 80 km) in isoprene-lined plastic boxes containing ice and frozen gel packs to the Laboratorio de Biología de Crustáceos, Centro Austral de Investigaciones Científicas (CADIC/CONICET) in Ushuaia where they were acclimatized, fully submerged for at least 5 days in 40-L plastic tanks containing running seawater.

Acclimatization was performed in a temperature-controlled room at 8 °C, under a 14 h light: 10 h dark photoperiod. The crabs were separated into groups of about 30 crabs each, according to species in tanks containing 30 L of running, recirculating seawater (5,000 L tank) at 7 °C and 30 ‰S, a salinity similar to that at the collecting sites, with pebbles for refuge.

The crabs were fed on small pieces of chopped squid in the morning every 3 days over the entire acclimatization and experimental period, except during the experiments to establish critical salinity limits, osmoregulatory capability and gene expression. Uneaten food fragments were removed in the evening of each feeding day. At the end of the experimental period, unused crabs were returned safely to their collection sites and released.

### Survival and estimation of critical salinity limits

After the 5-day acclimatization period, groups of intermolt crabs (stage C-D_0_, Drach and Tchernigovtzeff, 1967) of each species were assigned directly to aerated, covered plastic pots containing 3 L of seawater prepared at concentrations either above or below the acclimatization salinity of 30 ‰S, and held at 7 °C. Salinities above 30 ‰ S were prepared from the first thaw of frozen seawater (≈90 ‰S); those below were prepared with distilled water. All salinities were verified using an optical refractometer (American Optical Corp., Southbridge, MA, USA).

To establish the lower (LL_50_) and upper (UL_50_) 5-day critical salinity limits of 50% mortality for each species, groups of 7 crabs each were transferred directly to 2, 5 or 10, and 60 or 65 ‰S for *A. albatrossis*, and to 5, 10 or 20, and 50, 55 or 60 ‰S for *H. planatus*, respectively. Mortality was checked every 12 h. Crabs that could not right themselves when placed upside down and that showed no movements of their antennae when touched gently with a fine wire thread were considered ‘dead’.

The species’ critical limits were calculated using Probit analyses that adjust percentage survival to a linear regression model (Finney, 1971; Thurman, 2002, 2003; Maraschi et al. 2021). The LL_50_ and UL_50_ values were 4 and 63 ‰S for *A. albatrossis*, and 12 and 50 ‰S for *H. planatus*, respectively.

### Experimental design and the time course of salinity challenge

To establish osmotic and Na^+^ and Cl^−^ regulatory capabilities, tissue hydration levels and gill ion transporter gene expression for each species, groups of 7 crabs each were acclimated directly for 5 days (120 h) to salinities of 5, 10, 20, 30, 40, 50 or 60 ‰S for *A. albatrossis* and 10, 20, 30, 40 or 50 ‰S for *H. planatus* as described above.

To accompany the time courses of osmotic and ionic regulation and gill gene expression during hypo- or hyper-osmotic challenge, salinities corresponding to 80% of the LL_50_ (80% LL_50_) and UL_50_ (80% UL_50_) values (i. e., 5 and 50 ‰S for *A. albatrossis*, and 15 and 40 ‰S for *H. planatus*) were used, respectively. Thus, each species was challenged with equivalent, severe and symmetrical, but non-lethal salinities, enabling direct comparison of their responses (Mantovani and McNamara, 2021; Maraschi et al., 2021). Groups of 7 crabs each were exposed directly for 0 [=30 ‰S, control], 6, 24 or 120 h at these salinities, as given above.

After all exposure periods, the crabs were cryoanesthetized in crushed ice for 5 min to enable sampling of the hemolymph for osmotic and ionic analyses, of the abdominal and chela muscles to accompany tissue hydration, and to harvest the posterior gills for ion transporter gene sequencing and quantitative expression.

### Measurement of hemolymph osmolality and sodium and chloride concentrations

Individual hemolymph samples of 10 to 50 μL in volume were obtained from each crab’s ventral sinuses using a #25-7 gauge needle coupled to a 1-mL plastic syringe, inserted into the arthrodial membrane at the junctions of the pereiopods and the carapace. Samples were frozen in 200-μL Eppendorf microtubes at −20 °C for air transport in dry ice to the Laboratory of Crustacean Physiology in Brazil and later measurement.

After thawing and vortexing, the osmolality (mOsm kg^−1^ H_2_O) of each sample was measured in undiluted 10-μL aliquots using a vapor pressure micro-osmometer (Wescor, model 5500, Logan UT, USA). Chloride concentration (mmol L^−1^) was measured in undiluted 10-μL aliquots of the same samples by microtitration against mercury nitrate using s-diphenylcarbazone as an indicator, employing a microtitrator (Metrohm model E 485, Herisau, Switzerland) according to Schales and Schales (1941) adapted by Santos and McNamara (1996). Hemolymph Na^+^ concentration (mmol L^−1^) was measured by atomic absorption spectroscopy (GBC, model 932AA, GBC Scientific Equipment Ltd, Braeside, VIC, Australia) in 10-μL aliquots diluted 1: 25,000 with distilled water.

### Estimation of hemolymph osmotic, sodium and chloride regulatory capabilities

To evaluate osmotic, sodium and chloride regulatory capabilities, hemolymph osmolalities and sodium and chloride concentrations of the 5-day salinity acclimated crabs were fitted to third order polynomial equations using the curve fitting function of SlideWrite Plus for Windows 7 software (Advanced Graphics Software, Inc., Encinitas, CA, USA). The isosmotic, iso-sodium and iso-chloride points are the intercepts of the fitted curves with the respective isosmotic/iso-sodium/iso-chloride lines, each estimated using the curve fit data display function.

Hemolymph osmotic, sodium and chloride hyper- and hypo-regulatory capabilities were expressed numerically as the respective change in the hemolymph parameter concentration as a function of that in the external medium (Δ hemolymph parameter/Δ external medium parameter), above or below the isosmotic, iso-sodium or iso-chloride points, respectively. Ratios close to ‘1’ reveal little regulatory capability while values near ‘0’ indicate excellent regulatory capability.

### Muscle tissue hydration

After hemolymph sampling, each crab was killed by bisecting the cerebral and abdominal nerve ganglia, and a ≈70-mg sample of muscle tissue was dissected from the abdomen and chelae. The muscle fragments were placed in previously weighed Eppendorf microtubes (M_T_) and weighed immediately on an electronic analytical balance (Ohaus Analytical Plus AP250D, Parsippany, NJ, USA, ±10 μg precision), providing the tube and sample wet mass (M_T_ + M_W_). The tubes were then placed open in a drying oven at 60 °C for 24 h. The tubes and dried muscle fragments were then allowed to cool in a desiccator and were reweighed providing the tube and sample dry mass (M_T_ + M_D_). The tube weight was then subtracted from each sample (M_T_ - M_W_ and M_T_ - M_D_) and the degree of muscle tissue hydration (T_H_, in %) calculated as T_H_ = [(M_W_ - M_D_)/M_W_] x 100.

### RNA extraction and amplification of gill ion transporter partial cDNA sequences

The branchiostegites were removed from the freshly killed crabs and the three posterior gill pairs were dissected and set aside in TRIzol reagent (Life Technologies, Thermo Fisher Scientific, Waltham, MA, USA) for total RNA extraction. The molecular methods followed those set out in Mantovani and McNamara (2021) and Maraschi et al. (2021). Briefly, total RNA was extracted from the pooled 3 posterior gill pairs (≈50 mg) of each crab at each salinity and time combination in TRIzol (1: 10 w/v) under RNAse-free conditions. Individual samples were homogenized in a standard manner in Eppendorf microtubes for 20 s using a disposable polypropylene pellet pestle attached to a DeWalt DWD024K electric drill mounted on a laboratory stand. Extraction was interrupted at the 70% ethanol step when the samples were frozen at −80 °C for posterior air transport in dry ice and storage at −80 °C in the Laboratory of Crustacean Physiology in Brazil.

After thawing and further processing, the extracted total RNA was quantified (Qubit 2.0 fluorometer, Thermo Fisher Scientific, Waltham, MA, USA) and 1 μg total RNA was treated with RNAse-free DNAse I (Invitrogen). Reverse transcription of mRNA to cDNA was then performed using oligo(dT) primers and a Superscript III reverse transcriptase kit (Invitrogen), employing a Veriti thermal cycler (Thermo Fisher), according to the manufacturer’s instructions. Success of both the DNase I treatment and of the cDNA obtained was verified in all samples by PCR amplification of the partial coding region for ribosomal protein L10 (RPL10) using the appropriate standard primers (Table 1), and visualized in 1% agarose gels.

**Table 1.**
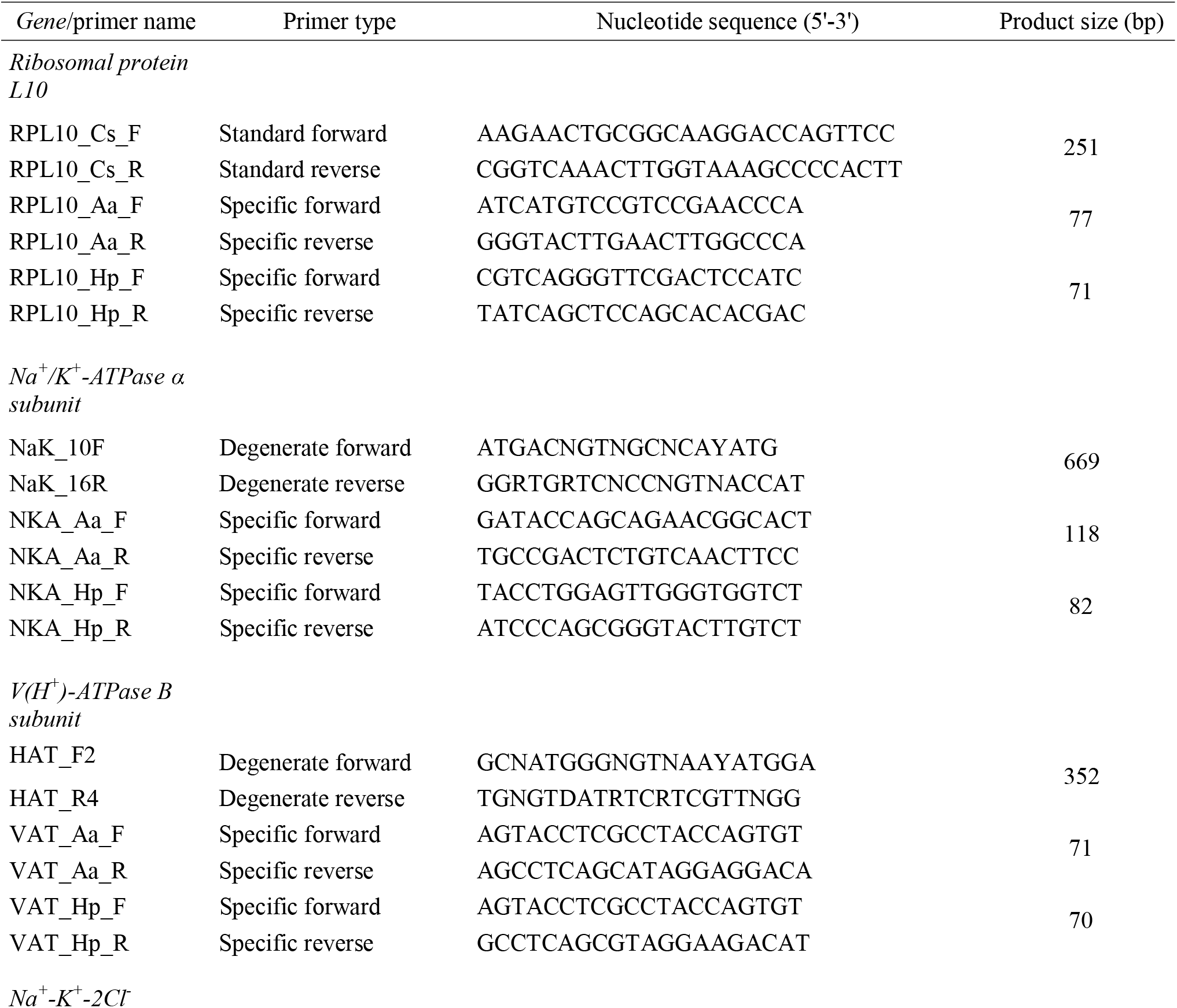
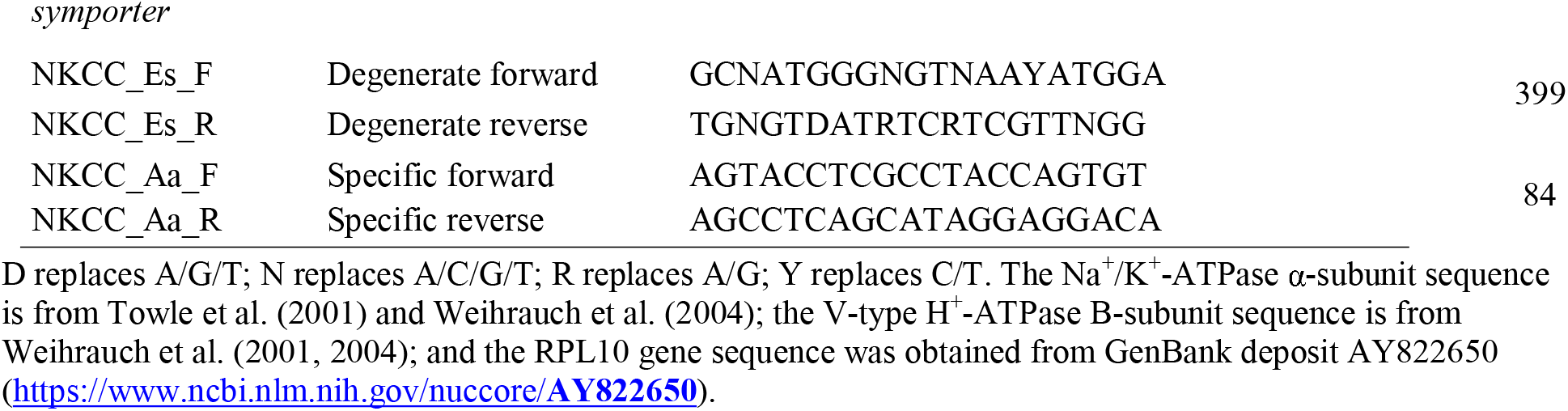
Standard, degenerate and specific oligonucleotide primers used to amplify partial coding regions of the genes for Ribosomal Protein L10 (RPL10_Cs_F and RPL10_Cs_R, RPL10_Aa_F and RPL10_Aa_R, RPL10_Hp_F and RPL10_Hp_R), Na^+^/K^+^-ATPase α-subunit (NaK_10F and NaK_16R, NKA_Aa_F and NKA_Aa_R, NKA_Hp_F and NKA_Hp_R), V-type H^+^-ATPase B-subunit (HAT_F2 and HAT_R4, VAT_Aa_F and VAT_Aa_R, VAT_Hp_F and VAT_Hp_R) and Na^+^/K^+^/2Cl^−^symporter (NKCC_Es_F and NKCC_Es_R, NKCC_Aa_F and NKCC_Aa_R) in the posterior gills of the Neotropical sub-Antarctic crabs Acanthocyclus albatrossis and Halicarcinus planatus.

### Cloning and sequencing of the partial cDNA sequences

The amplified fragments were cut out of their gel bands and purified using a PureLink Quick Gel Extract Kit (Thermo Fisher), cloned into a plasmid pCR 2.1-TOPO TA vector (Thermo Fisher) and transformed in thermocompetent DH5α *Escherichia coli*. Success of the transformation and choice of the recombinant plasmids was verified in LB agar cultures (Lennox L Agar, Thermo Fisher) containing Ampicillin (200 mg/mL) (Sigma-Aldrich, Missouri, EUA), X-Gal and IPTG (Thermo Fisher), followed by culture in an LB broth base (Thermo Fisher) containing Ampicillin (200 mg/mL). A diagnostic PCR was then run with the appropriate degenerate or standard primers (Table 1), with visualization in 1% agarose gels. The plasmids were then extracted and purified using a PureLink Plasmid Mini Kit (Thermo Fisher) and sequenced (Genetic Analyzer, ABI PRISM Model 3100, Applied Biosystems, Foster City, CA, USA) employing the dideoxinucleotide method (Sanger et al., 1977) using the primers (M13F e M13R) supplied with the vector kit.

After sequencing the amplified clones, the fragment sequences were analyzed for open reading frames (ORF). Searches of GenBank using the BLAST algorithm (Altschul et al., 1990)(http://www.ncbi.nih.gov/BLAST/) revealed high similarities of the nucleotide and predicted amino acid sequences of the target genes with sequences previously deposited for the coding regions analyzed in other crustacean species. The partial cDNA sequences obtained in the posterior gills for the RPL10 (GenBank accession number **<u>MG212501</u>**), Na^+^/K^+^-ATPase α-subunit (**<u>MG182147</u>**), V-ATPase B-subunit (**<u>MG212500</u>**) and Na^+^/K^+^/2Cl^−^ symporter (**<u>KM364038</u>**) genes in *A. albatrossis* posterior gills, and for the RPL10 (**<u>KM360152</u>**), Na^+^/K^+^-ATPase α-subunit (**<u>KM364036</u>**) and V-ATPase B-subunit (**<u>KM364037</u>**) genes in *H. planatus* were then employed to design specific primers for real-time quantitative gene expression (Primer-Blast, Primer Analysis Software, copyright 1989-91 Wojciech Ruchlik) (http://www.ncbi.nlm.nih.gov/tools/primer-blast/). Most unhappily, we were unable to clone the Na^+^/K^+^/2Cl^−^ symporter gene in *H. planatus*.

### Quantitative expression of gill ion transporter genes

The relative abundance of target gene mRNA in the total RNA extracts was estimated by quantitative reverse transcription (RT) real-time PCR (BioRad model CFX Connect, Hercules, CA, USA). Real-time PCR reactions (qPCR) were performed in triplicate using the *Power* SYBR Green PCR Master Mix Kit (Thermo Fisher) according to the manufacturer’s instructions, employing the specific primer pairs described in Table 1. Negative controls were performed without cDNA (‘No Template Control’) to detect eventual contamination.

The thermocycling procedure entailed an initial step at 95 °C for 10 min followed by 40 cycles of 15 s each at 95 °C and a final step at 60 °C for 1 min. The RPL10 gene that encodes for ribosomal protein L10 was used as an endogenous control. At the end of the reaction a dissociation curve was performed to verify eventual contamination, primer dimer formation or amplification of more than one amplicon.

Similarity between the amplification efficiencies [E=10^(−1/slope)^] of the target ion transporter genes and the endogenous RPL10 control gene for each species was evaluated by performing standard curve validations for all qPCR primers. Efficiencies were between 90% and 110%, with R^2^ >0.99. Relative mRNA expression of the Na^+^/K^+^-ATPase α-subunit, V-ATPase B-subunit and Na^+^/K^+^/2Cl^−^ symporter was normalized by the expression of the respective ribosomal protein L10 mRNA in the same sample for each species and condition.

To compare target gene expression after 120 h direct salinity challenge, and during the time course of hyper- or hypo-osmotic challenge (5 or 50 ‰S for *A. albatrossis*, 15 or 40 ‰S for *H. planatus*), the normalized data were calibrated by the respective mean gene expression for the control group (30 ‰S, time = 0 h), whose relative arbitrary expression was considered to be ‘1’. The calibrated data were treated using the exponential formula 2^−ΔΔCt^ (Livak and Schmittgen, 2001) and are given as the mean ± s.e.m.

The gill RPL10 gene was used as an endogenous control since it is expressed at very similar levels in various crustaceans held at different salinities and exposure intervals (Faleiros et al., 2010; Leone et al., 2015; Maraschi et al., 2021) and between different gills (Leone et al., 2015) and ammonia concentrations (Pinto et al., 2016). To illustrate, mean RPL10 Ct expression ranges from 1.0% to 2.5% and from 0.4% to 4.0%, in the shrimps *Palaemon northropi* and *Macrobrachium acanthurus*, respectively (Faleiros et al., 2017). In the freshwater crab *Dilocarcinus pagei* and shrimp *M. jelskii*, total Ct variation was 1.85 cycles for 15 salinity/exposure combinations, and 1.33 cycles for 12 salinity/exposure combinations, respectively (Mantovani and McNamara, 2021).

### Statistical analyses

After verifying normality of distribution (Kolmogorov-Smirnov or Shapiro-Wilk tests) and equality of variance (Levene’s or Brown-Forsythe test), the data were analyzed using one-way (salinity or exposure time) or two-way (species and salinity) analyses of variance to evaluate the main and interactive effects on hemolymph osmolality, chloride and sodium concentrations, on tissue hydration levels, and on ion transporter gene expression. Occasionally, the data were log transformed to meet the criteria for equal variance. The Student-Newman-Keuls *post-hoc* multiple means procedure was performed to locate statistically different means.

All analyses were performed using SigmaStat 2.03 (Systat Software Inc., San Jose, CA, USA), employing a minimum significance level of α= 0.05 with P-values ≤0.05 being considered significantly different. Data are expressed throughout as the Mean ± Standard Error of Mean and were plotted using SlideWrite Plus for Windows 7 software (Advanced Graphics Software, Inc., Encinitas, CA, USA).

## Results

### 1. Lower and upper critical salinity limits

*Acanthocyclus albatrossis* tolerates a slightly wider experimental salinity range (2-65 ‰S) than does *H. planatus* (5-60 ‰S). Their respective 5-day LL_50_ and UL_50_ critical salinities were 4 and 63 ‰S for *A. albatrossis*, and 12 and 50 ‰S for *H. planatus*.

### 2. Hemolymph osmotic and ionic regulation

#### Salinity challenge and regulatory capability

##### A. Osmolality

Two-way analysis of variance revealed marked effects of ‘salinity’ (F_4,59_ = 1524.2, P< 0.001), ‘species’ (F_1,59_= 64.1, P < 0.001) and their interaction (F_4,59_ = 38.3, P < 0.001) on hemolymph osmolality. Challenge at salinities of from 5 to 60 ‰S for 5 days revealed that *A. albatrossis* is a weak hyperosmotic regulator below 40 ‰S although tending to iso/hyper-conform at higher salinities (Fig. 2A). In contrast, over the range from 10 to 50 ‰S, *H. planatus* tended to slightly hyper-osmoconform, more evidently at 10-20 ‰S (Fig. 2A). The calculated isosmotic points were 1,380 mOsm kg^−1^ H_2_O (46.0 ‰S) for *A. albatrossis* and ≈1,340 mOsm kg^−1^ H_2_O (44.7 ‰S) for *H. planatus*, both crabs showing weak to poor hyper-regulatory capabilities (Δ_hemolymph osmolality_/Δ_medium osmolality_) of 0.70 and 0.79 respectively.

**Figure 2.**
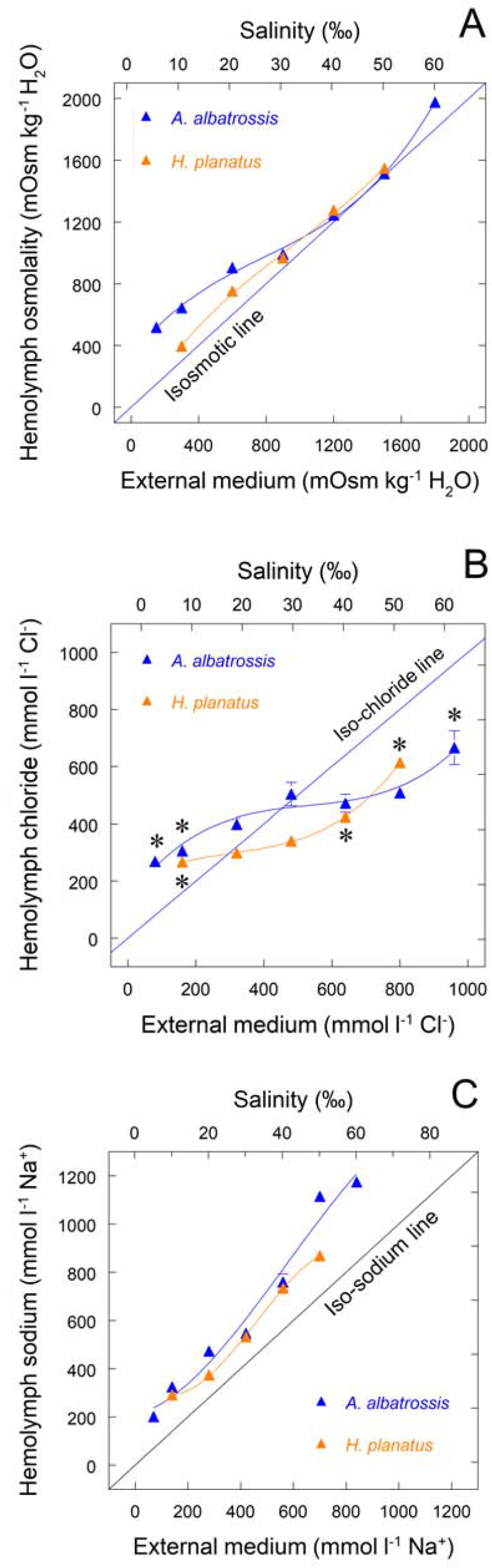
Hemolymph osmotic, chloride and sodium regulatory capability in salinity-acclimated sub-Antarctic crabs, *Acanthocyclus albatrossis* and *Halicarcinus planatus*. Crabs held at 7 °C were acclimated for 5 days by direct transfer from seawater (30 ‰S, control) to the selected salinities (5 to 60 ‰S for *A. albatrossis*, 10 to 50 ‰S for *H. planatus*). (A) *A. albatrossis* is a weak hyperosmotic regulator while *H. planatus* tends to osmoconform. (B) Both crabs hyper/hypo-regulate [Cl^−^] well, while (C) [Na^+^] is hyper-regulated at all salinities. The isosmotic and iso-chloride points were 1,380 mOsm kg^−1^ H_2_O and 452 mmol L^−1^ Cl^−^ for *A. albatrossis*, and ≈1,340 mOsm kg^−1^ H_2_O and 316 mmol L^−1^ Cl^−^ for *H. planatus* (1 ‰S = 30 mOsm kg^−1^ H_2_O, 16 mmol L^−1^ Cl^−^ and 14 mmol L^−1^ Na^+^). The lowest [Na^+^] calculated closest to the iso-sodium line were 356 mmol L^−1^ Na^+^ for *A. albatrossis* and 390 mmol L^−1^ Na^+^ for *H. planatus*. Data are the mean ± SEM (N = 7) and have been fitted to third order polynomial equations. Where lacking, error bars are smaller than the symbols used. For osmolality, all means were significantly different (ANOVA, SNK, P ≤ 0.05); for [Cl^−^], *P ≤ 0.05 compared to control crabs acclimatized at 30 ‰S; for [Na^+^], all means were significantly different from control crabs (P ≤ 0.05) except at 20 ‰S.

##### B. Chloride

‘Salinity’ (F_4,60_ = 28.5, P < 0.001) and the interaction between ‘species’ and ‘salinity’ (F_4,60_ = 6.8, P < 0.001), but not ‘species’ (F_1,60_ = 2.5, P < 0.1), affected hemolymph [Cl^−^]. Differently from osmolality, however, both species hyper/hypo-regulated hemolymph [Cl^−^] well over their respective salinity ranges after 5 days, although *A. albatrossis* exhibited a much better overall chloride capability than did *H. planatus* (Fig 2B). The iso-chloride points were 452 mmol L^−1^ Cl^−^ for *A. albatrossis* and 316 mmol L^−1^ Cl^−^ for *H. planatus*. Chloride hyper-regulatory abilities were strong at 0.49 and 0.21, respectively, while hypo-regulatory capabilities were 0.42 and 0.68, respectively.

##### C. Sodium

‘Salinity’ (F_4,60_ = 377.4, P < 0.001), ‘species’ (F_1,60_= 169.1, P < 0.001) and their interaction (F_4,60_ = 18.3, P < 0.001) all contributed significantly to variation in hemolymph [Na^+^]. In yet a different pattern, [Na^+^] was hyper-regulated at all salinities in both species, and particularly so in *A. albatrossis* (Fig. 2C) after 5-days challenge. The lowest hemolymph [Na^+^] calculated closest to the iso-sodium line were 356 mmol L^−1^ Na^+^ for *A. albatrossis* and 390 mmol L^−1^ Na^+^ for *H. planatus*.

#### Time course of challenge at 80%UL_50_ and 80%LL_50_ salinities

##### A. Osmolality

Two-way variance analysis revealed that ‘species’ (F_1,48_ = 543.3, P < 0.001), ‘exposure time’ (F_3,48_ = 426.1, P < 0.001) and their interaction (F_3,48_ = 55.6, P < 0.001) all contributed notably to variation in hemolymph osmolality at the 80%UL_50_ salinities (50 ‰S for *A. albatrossis* and 40 ‰S for *H. planatus*). At the 80%LL_50_ salinities (5 and 15 ‰S, respectively), ‘exposure time’ (F_3,45_ = 141.2, P < 0.001), ‘species’ (F_1,45_ = 6.9, P = 0.01) and the ‘species’ × ‘exposure time’ interaction (F_3,45_ < 15.6, P = 0.001) were likewise very significant.

The 5-day time course of change in hemolymph osmolality at salinity challenges corresponding to 80%UL_50_ and 80%LL_50_ (50 or 5 ‰S, respectively) in *A. albatrossis* showed that at 50 ‰S (1,500 mOsm kg^−1^ H_2_O), osmolality increased rapidly by 6 h, becoming isosmotic by 24 h, maintained up to 120 h (Fig. 3A). Asymmetrically, at 5 ‰S (150 mOsm kg^−1^ H_2_O), osmolality decreased gradually at 6 and 24 h, a gradient of Δ= +368 mOsm kg^−1^ H_2_O being maintained above ambient after 120 h exposure despite a further decrease (Fig. 3A).

**Figure 3.**
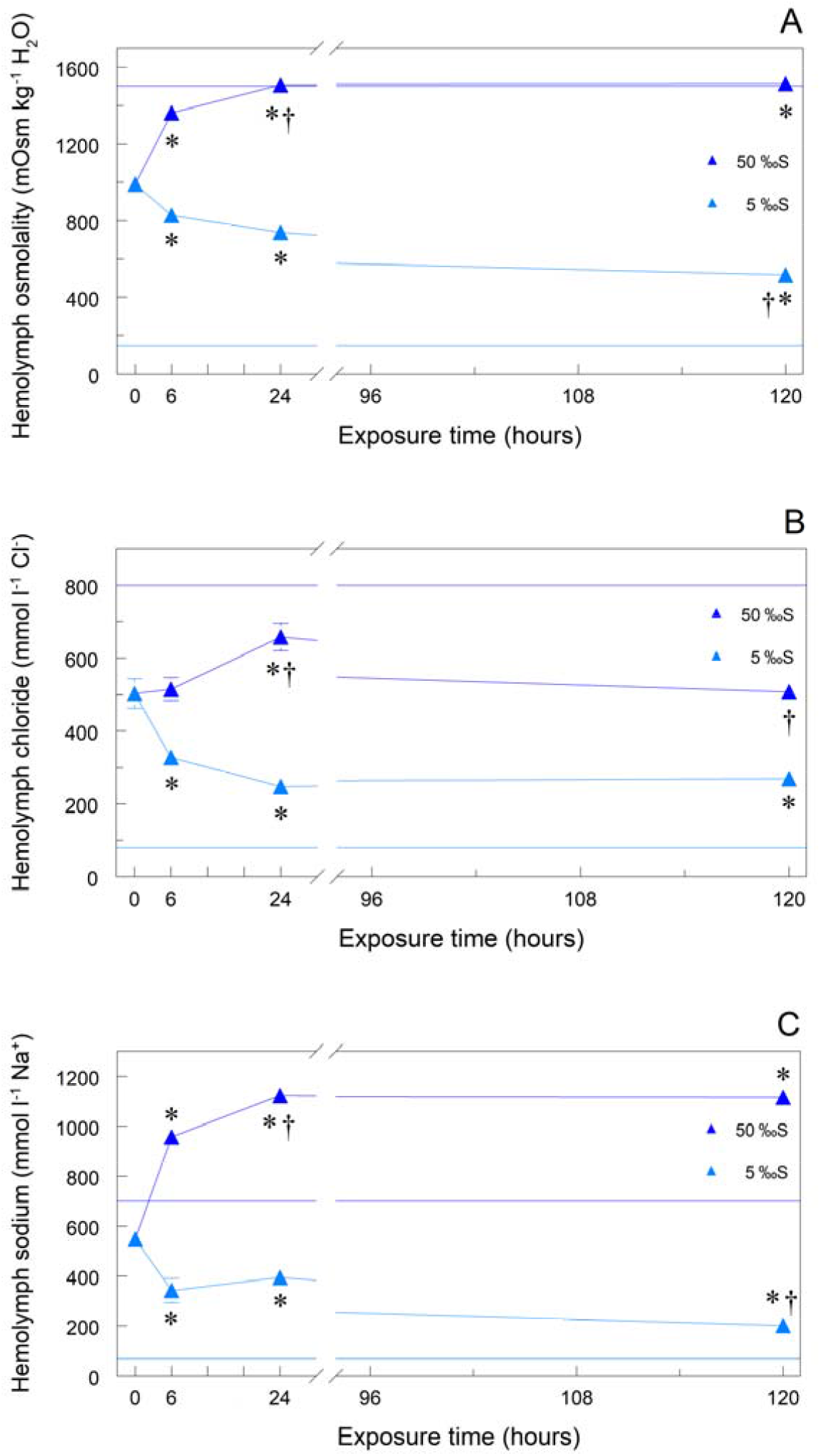
Time course of changes in osmoregulatory parameters in the sub-Antarctic crab *Acanthocyclus albatrossis* subjected to hyper-(50 ‰S, 80%UL_50_) or hypo-osmotic salinity (5 ‰S, 80%LL_50_) challenge for 5 days. (A) Hemolymph osmolality. (B) Hemolymph chloride concentration. (C) Hemolymph sodium concentration. Data are the mean ± SEM (N = 7). Where lacking, error bars are smaller than the symbols used. *P ≤ 0.05 compared to control crabs at Time = 0 h, †P ≤ 0.05 compared to immediately preceding value (ANOVA, SNK). Values for exposure at Time = 0 h are the respective hemolymph concentrations at 30 ‰S, the acclimatization salinity, at 7 °C. Upper and lower horizontal lines indicate the external medium osmolalities (1,500 and 150 mOsm kg^−1^ H_2_O), [Cl^−^] (800 and 80 mmol l^−1^) and [Na^+^] (700 and 70 mmol l^−1^) at the 80%UL_50_ and 80%LL_50_, respectively.

In *H. planatus* at 40 ‰S (80%UL_50_, 1,200 mOsm kg^−1^ H_2_O), hemolymph osmolality also increased rapidly at 6 and 24 h, becoming slightly hyper-osmotic after 120 h with Δ= +76 mOsm kg^−1^ H_2_O (Fig. 4A). At 15 ‰S (80%UL_50_, 450 mOsm kg^−1^ H_2_O), osmolality decreased very rapidly at 6 and 24 h, remaining stable and hyper-osmotic up to 120 h with Δ= +190 mOsm kg^−1^ H_2_O (Fig. 4A).

**Figure 4.**
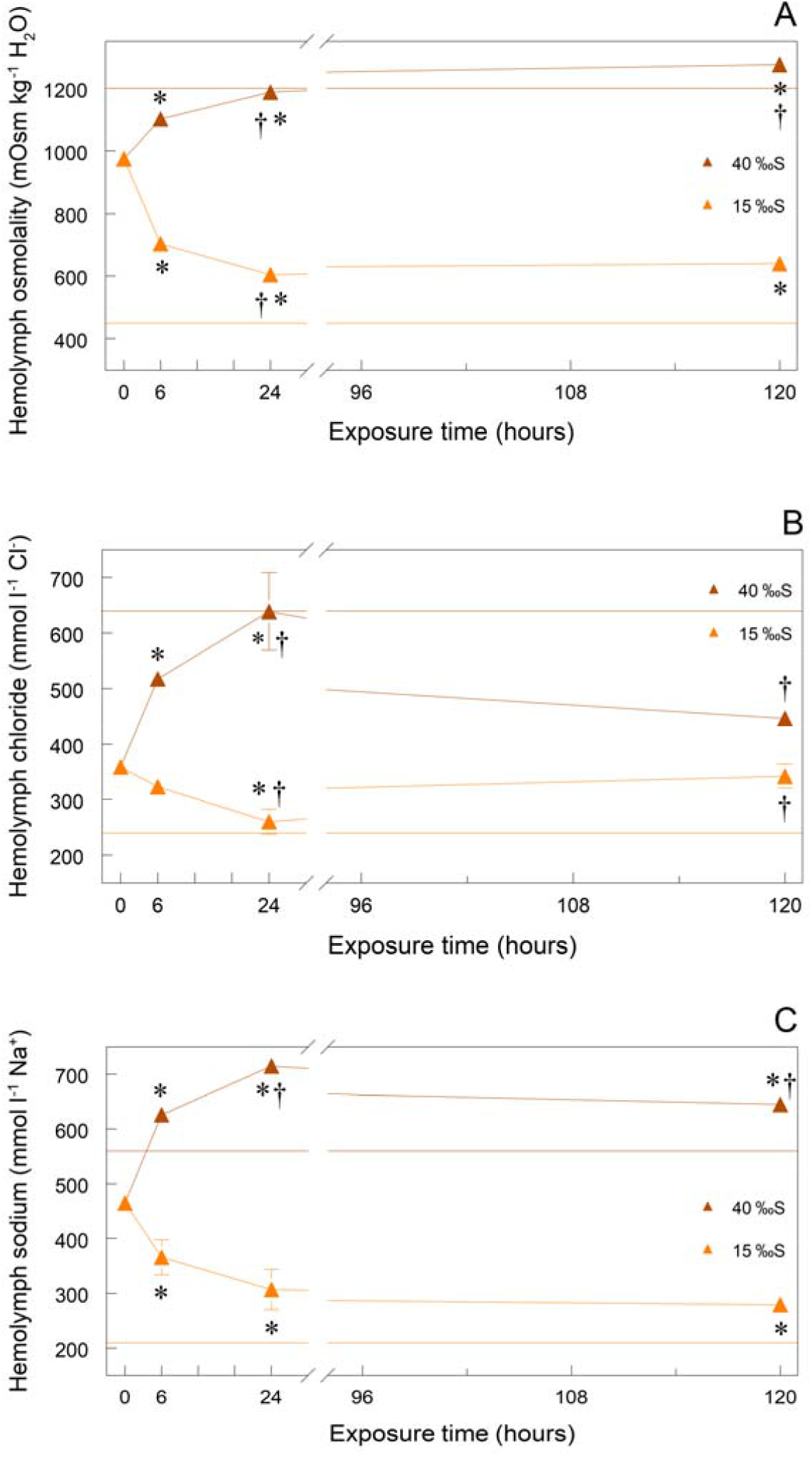
Time course of changes in osmoregulatory parameters in the sub-Antarctic crab *Halicarcinus planatus* subjected to hyper-(40 ‰S, 80%UL_50_) or hypo-osmotic salinity (15 ‰S, 80%LL_50_) challenge for 5 days. (A) Hemolymph osmolality. (B) Hemolymph chloride concentration. (C) Hemolymph sodium concentration. Data are the mean ± SEM (N = 7). Where lacking, error bars are smaller than the symbols used. *P ≤ 0.05 compared to control crabs at Time = 0 h, †P ≤ 0.05 compared to immediately preceding value (ANOVA, SNK). Values for exposure at Time = 0 h are the respective hemolymph concentrations at 30 ‰S, the acclimatization salinity, at 7 °C. Upper and lower horizontal lines indicate the external medium osmolalities (1,200 and 450 mOsm kg^−1^ H_2_O), [Cl^−^] (640 and 240 mmol l^−1^) and [Na^+^] (560 and 210 mmol l^−1^) at the 80%UL_50_ and 80%LL_50_, respectively.

##### B. Chloride

Two-way analysis of variance showed that ‘exposure time’ alone (F_3,48_ = 14.8, P < 0.001) affected hemolymph [Cl^−^] at the 80%UL_50_ salinities without ‘species’ or interactive effects. At the 80%LL_50_ salinities, ‘exposure time’ (F_3,48_ = 19.1, P < 0.001) and the interactive ‘species’ × ‘time’ (F_3,48_ = 7.0, P < 0.001) effects were significant factors.

Hemolymph [Cl^−^] in *A. albatrossis* was well regulated during the 5-day exposure time course (Fig. 3B). At 50 ‰S (800 mmol L^−1^ Cl^−^), [Cl^−^] increased at 24 h, returning to the outset value by 120 h, maintaining a gradient of Δ= −368 mmol L^−1^ Cl^−^ below ambient. At 5 ‰S (80 mmol L^−1^ Cl^−^), [Cl^−^] decreased progressively, remaining well above ambient from 24 h on with Δ= +190 mmol L^−1^ Cl^−^ at 120 h (Fig. 3B).

In *H. planatus* at 40 ‰S (640 mmol L^−1^ Cl^−^), hemolymph [Cl^−^] increased rapidly at 6, and at 24 h to iso-chloremic but becoming hypo-chloremic by 120 h with Δ= −194 mmol L^−1^ Cl^−^ below ambient (Fig. 4B). At 15 ‰S (240 mmol L^−1^ Cl^−^), [Cl^−^] decreased slowly to an iso-chloremic minimum at 24 h, increasing by 120 h with Δ= +102 mmol L^−1^ Cl^−^ above ambient (Fig. 4B).

##### C. Sodium

Two-way variance analysis revealed that ‘species’ (F_1,41_ = 458.4, P < 0.001), ‘exposure time’ (F_3,41_ = 152.3, P < 0.001) and their interaction (F_3,41_ = 31.7, P < 0.001) all markedly affected hemolymph [Na^+^] at the 80%UL_50_ salinities. At the 80%LL_50_ salinities, ‘exposure time’ (F_3,35_ = 35.6, P < 0.001) and the ‘species’ × ‘time’ interaction (F_3,35_ = 6.6, P = 0.001) were significant.

Hemolymph [Na^+^] in *A. albatrossis* at 50 ‰S (700 mmol L^−1^ Na^+^), increased rapidly above ambient at 6 and 24 h, remaining stable and hyper-natriuremic at 120 h with Δ= +415 mmol L^−1^ Na^+^ (Fig. 3C). At 5 ‰S (70 mmol L^−1^ Na^+^), hemolymph [Na^+^] declined rapidly at 6 h, continuing to a minimum above ambient Na^+^ at 120 h with Δ= +131 mmol L^−1^ Na^+^ (Fig. 3C).

In *H. planatus* at 40 ‰S (560 mmol L^−1^ Na^+^), hemolymph [Na^+^] increased rapidly at 6 and 24 h to well above ambient, decreasing slightly but remaining hyper-natriuremic by 120 h with Δ= +85 mmol L^−1^ Na^+^ (Fig. 4C). At 15 ‰S (210 mmol L^−1^ Na^+^), [Na^+^] decreased rapidly at 6 and 24 h, remaining unchanged although above ambient at 120 h with Δ= +69 mmol L^−1^ Na^+^ (Fig. 4C).

### 3. Muscle tissue hydration

Two-way variance analysis revealed a clear effect of ‘salinity’ (F_4,43_= 2.9, P= 0.03) and a marginal effect of ‘species’ (F_1,43_= 3.6, P= 0.06) on chela and abdominal muscle tissue hydration. After 5 days acclimation, the muscle tissue of both crab species clearly became hydrated (74-77%) at low salinities and dehydrated (64%) at high salinities compared to their respective controls at 30 ‰S (58-67%) (Fig. 5). Movement was asymmetrical, water gain at salinities below the controls (30 ‰S) being greater than loss at salinities above. Overall, *H. planatus* was nominally more hydrated (Δ= +5%) than *A. albatrossis* at the same salinity.

**Figure 5.**
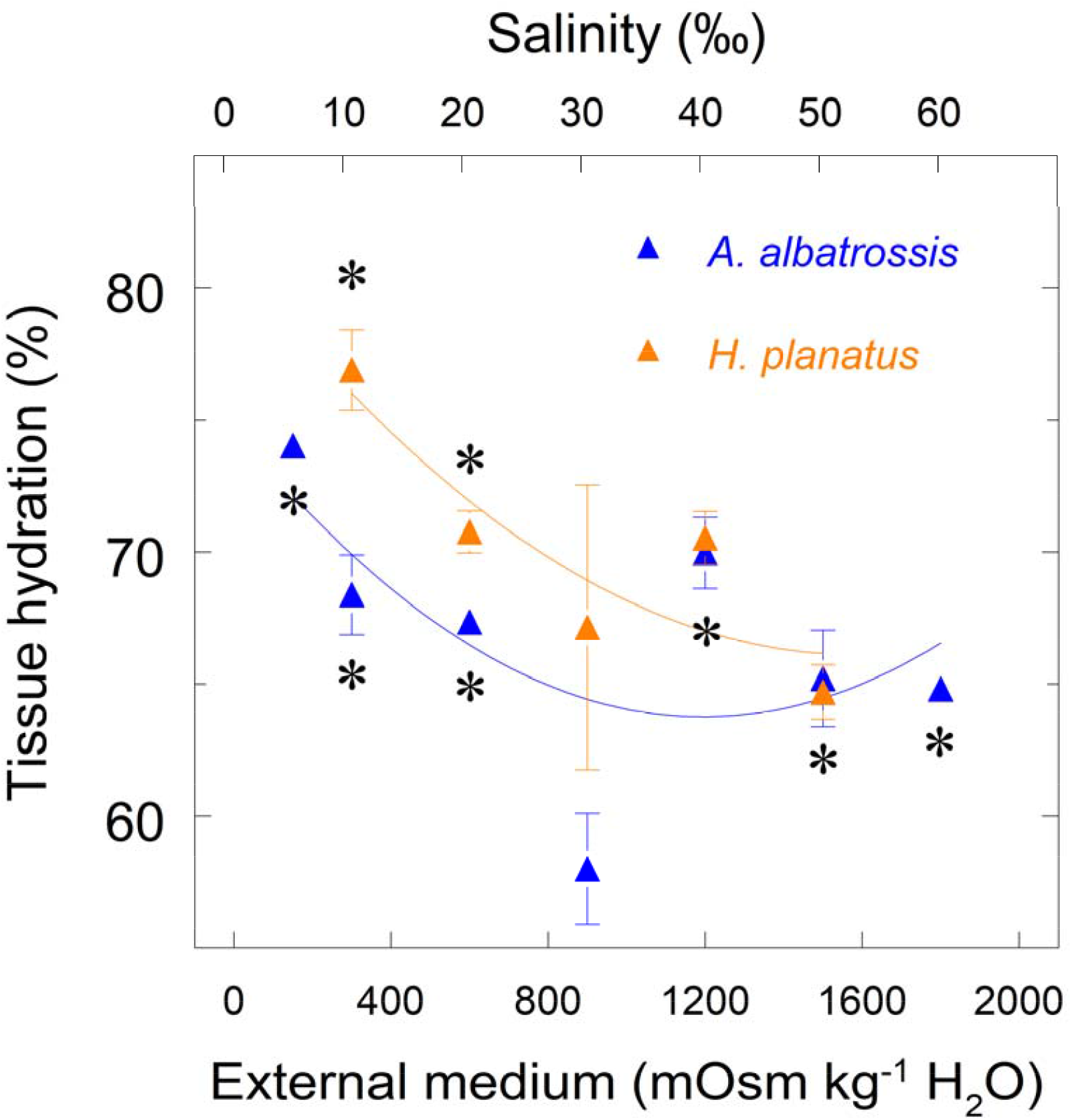
Muscle tissue hydration levels in salinity-acclimated sub-Antarctic crabs, *Acanthocyclus albatrossis* and *Halicarcinus planatus*. Crabs held at 7 °C were acclimated for 5 days by direct transfer from seawater (30 ‰S, control) to the selected salinities (5 to 60 ‰S for *A. albatrossis*, 10 to 50 ‰S for *H. planatus*). Tissue hydration increases at lower salinities and decreases at higher salinities in both species. Data are the mean ± SEM (N = 7) and have been fitted to quadratic equations. Where lacking, error bars are smaller than the symbols used. *P ≤ 0.05 compared to control crabs acclimated at 30 ‰S. †P ≤ 0.05 compared to crabs acclimated at 50 ‰S (ANOVA, SNK).

### 4. Gill ion transporter gene expression

#### Salinity and mRNA expression

##### A. Na^+^/K^+^-ATPase α-subunit

Two-way variance analysis revealed strong effects of ‘salinity’ (F_4,43_ = 13.2, P < 0.001), ‘species’ (F_1,43_= 14.2, P< 0.001) and their interaction (F_4,43_ = 2.9, P < 0.001) on α-subunit expression (Fig. 6A).

**Figure 6.**
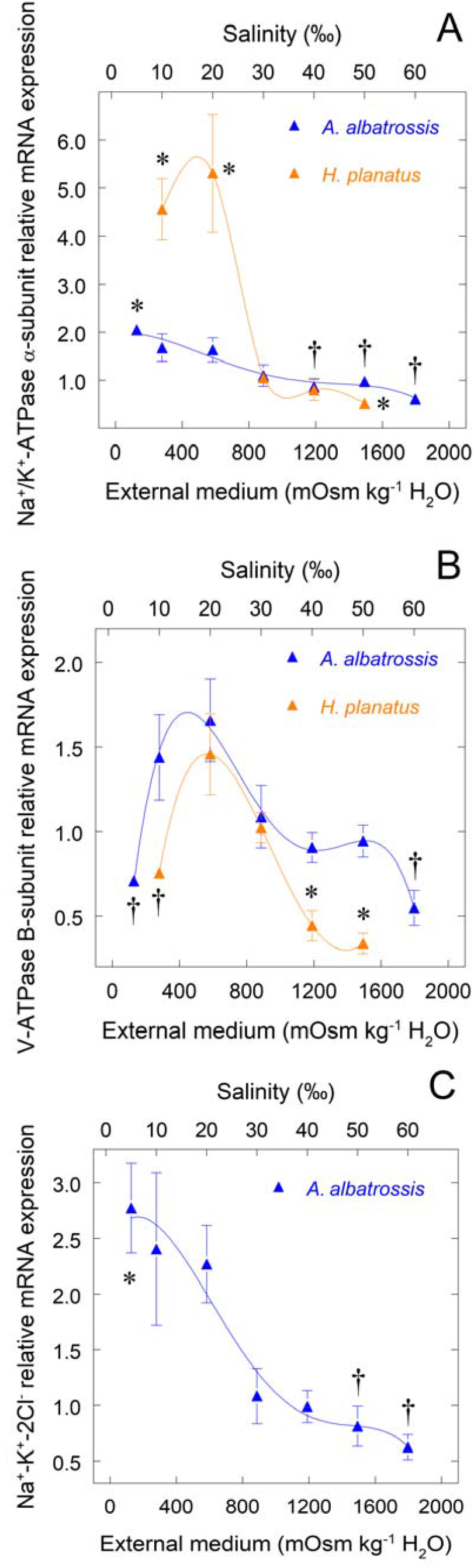
Relative mRNA expression of posterior gill ion transporter genes in salinity-acclimated sub-Antarctic crabs, *Acanthocyclus albatrossis* and *Halicarcinus planatus*. Crabs held at 7 °C were acclimated for 5 days by direct transfer from seawater (30 ‰S, control) to the selected salinities (5 to 60 ‰S for *A. albatrossis*, 10 to 50 ‰S for *H. planatus*); the three posterior gill pairs from individual crabs of each species were sampled. Expressions were calculated using the comparative C_t_ method (2^−ΔΔCT^) and have been normalized by expression of the constitutive RPL10 gene in the same sample and calibrated against the control expression at 30 ‰S. Data are the mean ± SEM (N = 7) and have been fitted to fourth order polynomial equations. Where lacking, error bars are smaller than the symbols used. (A) Na^+^/K^+^-ATPase α-subunit, *P ≤ 0.05 compared to control crabs acclimatized at 30 ‰S, †P < 0.05 compared to 5 ‰S. (B) V(H^+^)-ATPase B-subunit, *P ≤ 0.05 compared to control crabs acclimatized at 30 ‰S, †P < 0.05 compared to 20 ‰S. (C) Na^+^-K^+^-2Cl^−^ symporter, *P < 0.05 compared to control crabs acclimatized at 30 ‰S, †P < 0.05 compared to 20 ‰S (ANOVA, SNK). The Na^+^-K^+^-2Cl^−^ symporter could not be cloned in *H. planatus*.

##### B. V(H^+^)-ATPase B-subunit

Two-way analysis of variance disclosed notable effects of ‘salinity’ (F_4,44_= 9.2, P< 0.001) and ‘species’ (F_1,44_= 11.4, P< 0.001) on B-subunit expression. mRNA expression of the gill V(H^+^)-ATPase in *A. albatrossis* decreased moderately at both low (5 ‰S, 0.7-fold, −35%) and high (60 ‰S, 0.5-fold, −50%) salinities after 120 h acclimation (Fig. 6B) compared to expression at 20 ‰S. Similarly, in *H. planatus*, V(H^+^)-ATPase mRNA expression also decreased at low (10 ‰S, 0.7-fold, −26%) and high (50 ‰S, 0.3-fold, −67%) salinities (Fig 6B), showing greater sensibility at the high salinity extreme.

##### C. Na^+^-K^+^-2Cl^−^ symporter

The gill NKCC gene could only be cloned in *A. albatrossis*. After 120 h acclimation, mRNA expression of the NKCC symporter gene increased notably at low salinities between 5 and 20 ‰S (2.6-fold at 5 ‰S) and decreased to a lesser extent at 60 ‰S (0.6-fold, −42%) over control expression at 30 ‰S (Fig. 6C). Expressions at salinities between 30 and 60 ‰S were similar.

#### Time course of challenge at 80%UL_50_ and 80%LL_50_ salinities

##### A. Na^+^/K^+^-ATPase α-subunit

Two-way variance analysis revealed that ‘exposure time’ (F_3,30_ = 3.2, P = 0.04) alone contributed to variation in α-subunit expression at the 80%UL_50_ salinities (50 ‰S for *A. albatrossis* and 40 ‰S for *H. planatus*) with no effect of ‘species’. At the 80%LL_50_ salinities (5 and 15 ‰S, respectively), ‘exposure time’ (F_3,30_ = 8.0, P < 0.001), and the ‘species’ × ‘time’ interaction (F_3,30_ = 3.5, P = 0.03) contributed significantly, again with no difference between species.

The 5-day time course of change in gill Na^+^/K^+^-ATPase mRNA expression at salinity challenges corresponding to 80%UL_50_ and 80%LL_50_ (50 or 5 ‰S, respectively) in *A. albatrossis* showed that expression at both salinities increased rapidly by 6 h and was sustained until 24 h (Fig. 7A). At 50 ‰S, expression declined to outset values while at 5 ‰S, expression increased 1.9-fold by 120 h. In *H. planatus* at both 40 and 15 ‰S, mRNA expression was unchanged up to 24 h, and was sustained at control values after 120 h at 40 ‰S (Fig. 8A). However, at 15 ‰S, expression increased markedly by 8.2-fold after 120 h (Fig. 8A).

**Figure 7.**
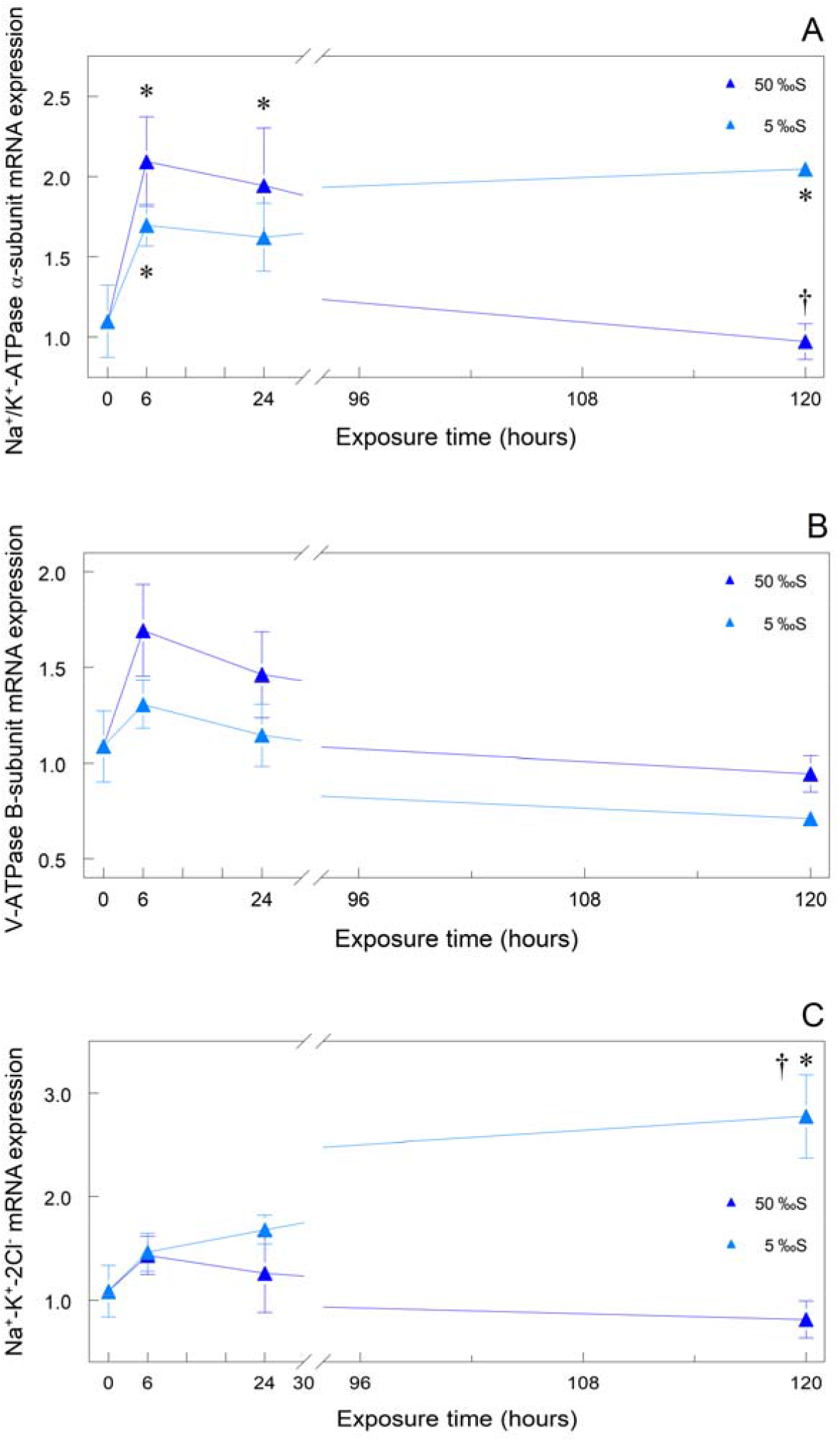
Time course of changes in relative mRNA expression of posterior gill ion transporter genes in salinity-acclimated sub-Antarctic crabs, *Acanthocyclus albatrossis* subjected to hyper-(50 ‰S, 80%UL_50_) or hypo-osmotic salinity (5 ‰S, 80%LL_50_) challenge for 5 days. (A) Na^+^/K^+^-ATPase α-subunit. (B) V(H^+^)-ATPase B-subunit. (C) Na^+^-K^+^-2Cl^−^ symporter. Data are the mean ± SEM (N = 7). Where lacking, error bars are smaller than the symbols used. *P ≤ 0.05 compared to control value, †P ≤ 0.05 compared to immediately preceding value (ANOVA, SNK). Values for exposure at Time = 0 h are the respective expressions at 30 ‰S, the acclimatization salinity, at 7 °C. Expressions were calculated using the comparative C_t_ method (2^−ΔΔCT^) and have been normalized by expression of the constitutive RPL10 gene in the same sample and calibrated against the control expression at Time = 0 h.

**Figure 8.**
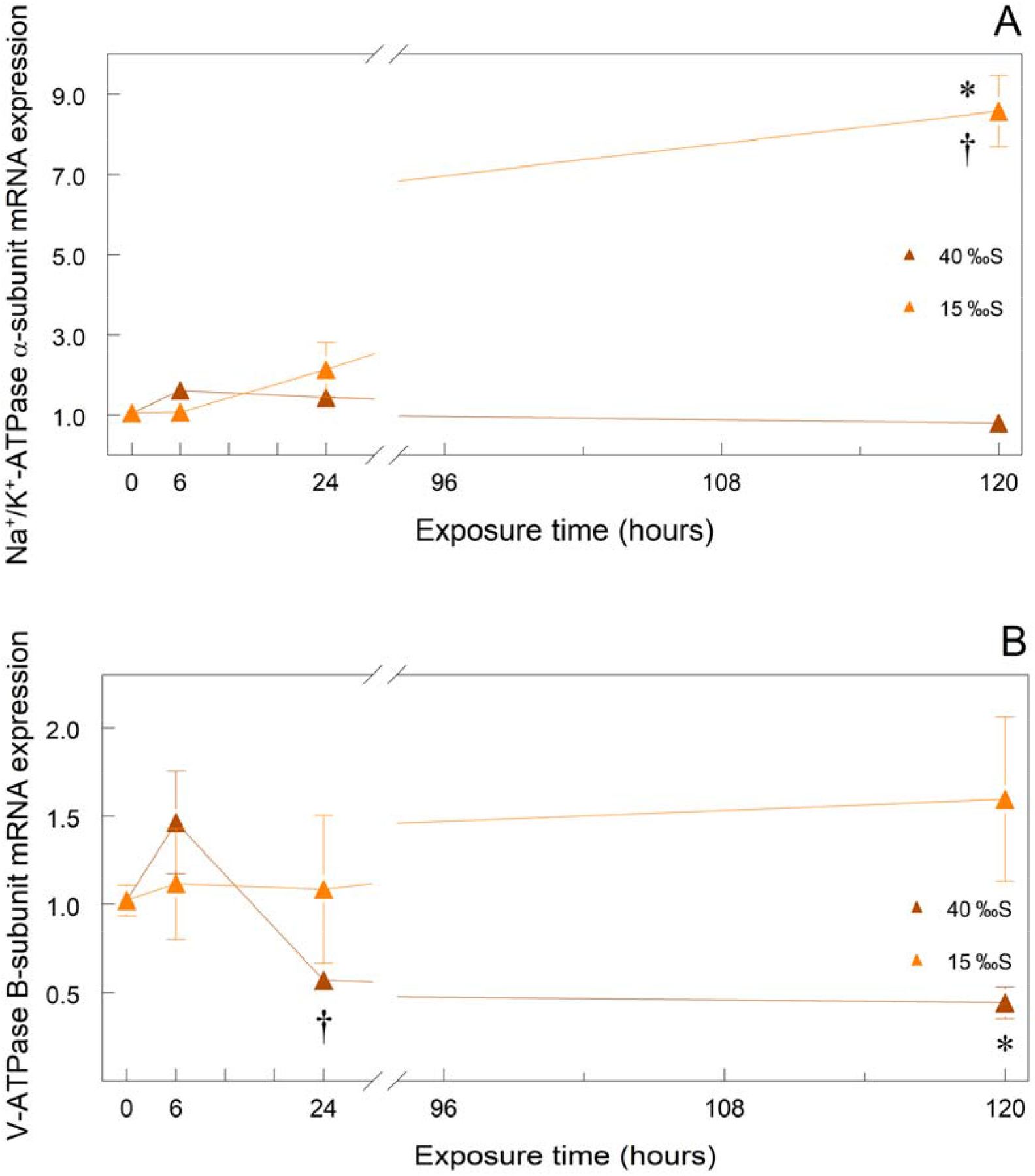
Time course of changes in relative mRNA expression of posterior gill ion transporter genes in salinity-acclimated sub-Antarctic crabs, *Halicarcinus planatus* subjected to hyper-(40 ‰S, 80%UL_50_) or hypo-osmotic salinity (15 ‰S, 80%LL_50_) challenge for 5 days. (A) Na^+^/K^+^-ATPase α-subunit. (B) V(H^+^)-ATPase B-subunit. Data are the mean ± SEM (N = 7). Where lacking, error bars are smaller than the symbols used. *P ≤ 0.05 compared to control value, †P ≤ 0.05 compared to immediately preceding value (ANOVA, SNK). Values for exposure at Time = 0 h are the respective expressions at 30 ‰S, the acclimatization salinity, at 7 °C. Expressions were calculated using the comparative C_t_ method (2^−ΔΔCT^) and have been normalized by expression of the constitutive RPL10 gene in the same sample and calibrated against the control expression at Time = 0 h. The Na^+^-K^+^-2Cl^−^ symporter could not be cloned.

##### B. V(H^+^)-ATPase B-subunit

Two-way analysis of variance revealed that ‘species’ (F_1,31_ = 5.7, P = 0.03) and ‘exposure time’ (F_3,31_ = 4.4, P = 0.01) contributed to variation in B-subunit expression at the 80%UL_50_ salinities (50 and 40 ‰, respectively). At the 80%LL_50_ salinities (5 and 15 ‰S, respectively), expression was not affected by any factor (0.8 < F_1/3,31_ < 2.3, 0.1 < P < 0.9).

Gill V(H^+^)-ATPase B-subunit mRNA expression was quantitatively unchanged in *A. albatrossis*. Despite a slight nominal increase (1.6-fold) by 6 h at 50 ‰S, V(H^+^)-ATPase mRNA expression in *A. albatrossis* showed a clear tendency to diminish thereafter to outset values over the 120-h time course (−44%, 1-way ANOVA F_3,15_= 3.164, P= 0.055) (Fig. 7B). A similar tendency to decrease was seen at 5 ‰S (−46%, 1-way ANOVA F_3,17_= 2.581, P= 0.087) (Fig. 7B). In *H. planatus* at 40 ‰S, gill V(H^+^)-ATPase B-subunit mRNA expression was unchanged although increased nominally by 1.4-fold at 6 h, subsequently declining 0.6-fold (−44%) by 24 h and sustained significantly below the control value at 120 h (−57%) (Fig. 8B). At 15 ‰S, expression was unchanged up to 120 h, although increasing nominally by 1.6-fold at 120 h (Fig. 8B).

##### C. Na^+^-K^+^-2Cl^−^ symporter

Data are available for *A. albatrossis* alone since we were unable to clone the gill NKCC gene in *H. planatus*. Gill NKCC mRNA expression in *A. albatrossis* was unchanged up to 24 h at both 50 and 5 ‰S, remaining at control levels at 50 ‰S (Fig. 7C). Expression at 5 ‰S increased notably by 2.6-fold at 120 h (Fig. 7C).

## Discussion

### Osmotic and ionic regulation

This is the first investigation of osmotic and ionic regulation and gill ion transporter gene expression in high latitude, Neotropical, subAntarctic crabs. We disclose that, despite sharing the same habitat and osmotic niche in the Beagle Channel, *Acanthocyclus albatrossis* and *Halicarcinus planatus* exhibit similar systemic hemolymph osmotic, Na^+^ and Cl^−^ regulatory abilities, but show clear differences in salinity tolerance, and in gill Na^+^/K^+^- and V(H^+^)-ATPase and Na^+^-K^+^-2Cl^−^ symporter mRNA expression on rigorous salinity challenge. Hemolymph [Cl^−^] is well regulated while [Na^+^] is not regulated. We analyze these species-specific divergences, evaluate adaptedness and seek to correlate relevant expression of ion transporter mRNA with the crabs’ abilities to effect regulatory salt uptake and secretion.

Both crabs exhibit very ample 5-day salinity tolerances, from around 2-5 to 60-65 ‰S, which are far greater than seen in many infralittoral crabs among the Portunidae, Cancridae and Varunidae. *Acanthocyclus albatrossis* tolerates an experimental salinity range about 14% wider than does *Halicarcinus planatus*, displaying greater lower and upper critical salinity limits. *Halicarcinus planatus* occurs in less variable salinities of around 30 ‰S (Benavides et al. 2019), and survives experimental acclimation to dilute media down to 18 ‰S, showing 50% mortality after 36 days, but not surviving at 5 or 11 ‰S for more than a few days (López-Farrán et al. 2021). Apparently, *A. albatrossis* is more tolerant of dilute media to which it may be better adapted. Its occupation of a habitat characterized by lower all round salinities (≈26 ‰S, Zangrando et al. 2016), owing to freshwater outflow from the Río Varela into Bahía Varela and the adjacent Río Cambaceres, where the species tends to predominate, may have driven adaptation to this lower salinity limit. The reasons for the extended upper salinity limits (65 cf. 60 ‰S) in both species are not clear but may be intrinsic to their evolutionary history since surface salinities in the Beagle Channel do not exceed 32.7 ‰S (Isla et al. 1999) and are unlikely to have driven such a high upper critical limit.

Osmotic and ionic regulatory capacities in the two species are fairly similar overall, despite a species effect on hemolymph osmolality and sodium concentration. However, their abilities to regulate osmolality and specific ions show striking differences. *Acanthocyclus albatrossis* is a weak hyper-osmoregulator in dilute media, but osmoconforms in seawater and at salinities above ≈40 ‰S, much like other infralittoral crabs such as the portunids *Carcinus maenas* (Siebers et al. 1982, Cieluch et al. 2004) and *Callinectes ornatus* (Garçon et al. 2009), cancrid *Cancer irroratus* (Cantelmo et al. 1975), varunids *Hemigrapsus oregonensis, H. nudus* (Dehnel 1962) and *H. sanguineus* (Hudson et al. 2018) and panopeid *Rhithropanopeus harrisii* (Smith 1967). In contrast, *Halicarcinus planatus* osmoconforms over the 10 to 50 ‰S range used, much like the cancrid *Cancer pagurus* (Whitely et al. 2018), epialtids *Libinia emarginata, L. dubia* (Bursey 1982) and *Pugettia producta* (Cornell 1979) and hepatid *Hepatus pudibundus* (Freire et al. 2008b).

Despite a difference between species, neither crab is able to regulate hemolymph Na^+^, which is hyper-regulated at mean values of ≈235 mmol L^−1^ and at ≈200 mmol L^−1^ above iso-natriuretic in *A. albatrossis* and *H. planatus*, respectively, the difference tending to increase even further at salinities above seawater. These unregulated, high Na^+^ concentrations found over the salinity ranges tested appear to underlie the species’ inability to regulate hemolymph osmolality. In contrast, both species strongly regulate hemolymph Cl^−^, although more so in *A. albatrossis* that maintains larger gradients near its limits of salinity tolerance than does *H. planatus*. The much higher iso-chloride point in *A. albatrossis* (452 mmol L^−1^ Cl^−^ cf. 316 mmol L^−1^ Cl^−^ for *H. planatus*) and its better overall Cl^−^ hypo-regulatory ability suggest adaptation to a more saline environment despite other findings revealing tolerance of low salinity.

Muscle tissue hydration in *H. planatus* responds fairly linearly to osmotic challenge, gaining or losing water, respectively, in dilute or concentrated media, indicating reliance on mechanisms of isosmotic intracellular regulation (IIR) and organic osmolytes to buffer the changes in cell volume inherent to crustaceans that use an osmoconforming strategy (Augusto et al. 2007; Freire et al. 2008b; Foster et al. 2010). Although *A. albatrossis* shows a similar response, the significant species effect reveals lower overall tissue hydration levels and a flatter response curve, indicating less dependence on IIR, as reflected in its strong ability to regulate hemolymph Cl^−^, weak hyper-regulatory capacity and wider experimental salinity tolerance. These findings reveal that the two crabs have evolved subtly different adaptive osmoregulatory mechanisms expressed at the systemic level.

The time courses of exposure to 80% upper (80%UL_50_) or lower critical limit salinities (80%LL_50_) reveal that, despite a clear species effect, within 24 h both crabs become isosmotic at 80%UL_50_. In contrast, hemolymph [Na^+^] rapidly (6 h) exceeds iso-natriuretic and remains elevated, particularly in *A. albatrossis* that, unlike *H. planatus*, shows no sign of regulation. Hemolymph osmolality and [Na^+^] are much better regulated at the 80%LL_50_, neither species becoming isosmotic or iso-natriuretic. Na^+^ is lost more slowly and to a lesser extent in *A. albatrossis*. In sharp contrast, hemolymph [Cl^−^] is temporally well regulated in both species, particularly in *A. albatrossis* that recovers initial [Cl^−^] after 5 days at the 80%UL_50_ although not at the 80%LL_50_. *Halicarcinus planatus* becomes iso-chloremic initially at the 80%UL_50_ but recovers [Cl^−^] fully, also seen at the 80%LL_50_. These findings suggest that both species are better adapted to dilute than to concentrated media, particularly *A. albatrossis* with regard to osmolality and Na^+^. Chloride is more tightly regulated in *A. albatrossis* in concentrated media while *H. planatus* recovers completely on both low and high salinity challenge, again revealing subtle interspecific differences in Cl^−^ regulatory ability.

### Expression of gill ion transporter genes

As seen in many osmoregulating crabs (Luquet et al. 2005; Jayasundara et al. 2007; Jilette et al. 2010; Chen et al. 2019), gill Na^+^/K^+^-ATPase α-subunit mRNA expression decreases at high salinities in both *A. albatrossis* and *H. planatus*. While this may diminish the active Na^+^/K^+^-ATPase-driven component of total Na^+^ influx across the gill epithelium into the hemolymph, expression remains at ≈50% compared to isosmotic crabs and, together with Na^+^ flowing through the Na^+^-K^+^-2Cl^−^ symporter (see below) and possibly the Na^+^/H^+^ exchanger, is likely responsible for the continually increasing Na^+^ gradient seen at high salinities in both species. Apparently, passive Na^+^ permeability does not diminish either and also contributes to the species’ remarkably elevated hemolymph [Na^+^] at high salinity. Partly underlying the clear species effect, this reduction in mRNA expression is more evident in *H. planatus* while expression in *A. albatrossis* seems to be Na^+^ insensitive.

In marked contrast, gill Na^+^/K^+^-ATPase mRNA expression at low salinities differs conspicuously between the two species, doubling in *A. albatrossis* but increasing ≈5-fold in *H. planatus*, as seen in many hyper-osmoregulating crabs (Wang et al. 2018; Chen et al. 2019). However, this substantial up-regulation does not result in increased hemolymph Na^+^ concentrations, which are just 2-to 3-fold above ambient compared to [Na^+^] in moderate hyper-osmoregulators like the blue crab *C. danae* (4-fold, Garçon et al. 2021), swamp ghost crab *Ucides cordatus* (6-fold, Leone et al. 2020), thin-striped hermit crab *Clibanarius symmetricus* (3.4-fold, Faleiros et al. 2018), and to strong hyper-regulators like the mudflat fiddler crab *Minuca rapax* (>300-fold, Capparelli et al. 2017). Apparently, augmented gill Na^+^/K^+^-ATPase mRNA expression and consequent enzyme transport activity drive other ion capture pathways such as Cl^−^ uptake, independently of Na^+^ transport, particularly in *H. planatus*.

Chloride uptake at low salinities in *A. albatrossis* appears to be mediated by an apically-located Na^+^-K^+^-2Cl^−^ symporter (McNamara and Faria 2012) since mRNA expression of this gene increases ≈2.5-fold compared to crabs at isosmotic and higher salinities. Curiously, in estuarine and freshwater palaemonid shrimps, the gill Na^+^-K^+^-2Cl^−^ symporter also appears to underlie hemolymph [Cl^−^] hyper-regulation and hyper-osmoregulation (Maraschi et al. 2021). An evolutionary trade-off may have become established between the putatively deleterious effects of hemolymph Na^+^ and Cl^−^ imbalance and poorly hyper-regulated [Na^+^] against strongly regulated [Cl^−^] at low salinities, since Cl^−^ influx via the Na^+^-K^+^-2Cl^−^ symporter would be driven by external Na^+^ down its small electrochemical gradient into the apical cytosol (Freire et al. 2008a; McNamara and Faria 2012), contributing to transepithelial Cl^−^ uptake. Thus, even the modest 2-fold increase in Na^+^/K^+^-ATPase α-subunit expression seen in *A. albatrossis* at low salinities may be sufficient to drive cytosolic Na^+^ into the hemolymph accompanied by Na^+^-K^+^-2Cl^−^-mediated transepithelial Cl^−^ influx.

The strong hemolymph Cl^−^ hypo-regulation seen in *A. albatrossis* at high salinities is not dependent on altered Na^+^-K^+^-2Cl^−^ expression, which remains similar to isosmotic crabs. The very elevated hemolymph [Na^+^] would drive cytosolic Cl^−^ influx through a basally-located Na^+^-K^+^-2Cl^−^ symporter (McNamara and Faria 2012), also independently of Na^+^/K^+^-ATPase mRNA expression that is unaltered. The trade-off above also may include advantageous use of the strong hemolymph: cytosol Na^+^ gradient to drive Cl^−^ secretion at high salinity, independently of mainly unaltered Na^+^-K^+^-2Cl^−^ and Na^+^/K^+^-ATPase expressions. Nevertheless, an as yet undisclosed mechanism of active Cl^−^ secretion warrants investigation (see Gerencser and Zhang 2003). Most unfortunately, despite much effort, we were unable to clone the Na^+^-K^+^-2Cl^−^ symporter in *H. planatus*.

Alterations in gill V(H^+^)-ATPase B-subunit expression are subtle and similar overall in both crabs, despite a species effect. Expression decreases at high salinities, particularly in *H. planatus* that shows lower transcription in general like some other crabs (Tsai and Lin, 2007; Firmino et al. 2011), possibly reflecting diminished Na^+^/H^+^ antiporter availability. The decline in expression at the lower critical limits is unexpected (see Luquet et al. 2005). The ≈1.5-fold increase in expression at moderately dilute salinities suggests a putative role in driving Na^+^ uptake via the Na^+^/H^+^ antiporter or protonation of NH_3_ to excretable and/or Na^+^-exchangable NH_4_^+^ (Weihrauch et al. 2017).

The time courses of alterations in ion transporter gene mRNA expression reveal few species-specific differences. In *A. albatrossis*, Na^+^/K^+^-ATPase α-subunit expression increases 2-fold initially at both the 80%UL_50_ and 80%LL_50_, possibly underlying an early (6 to 24 h) Cl^−^ secretion mechanism, and Cl^−^ uptake in dilute medium putatively via an apical Cl^−^/HCO_3_ ^−^ antiporter. This increase is sustained at the 80%LL_50_, suggesting that increased Na^+^/K^+^-ATPase activity indirectly drives Cl^−^ uptake, but returns to outset expression levels at the 80%UL_50_, consistent with diminished overall Na^+^/K^+^-ATPase-dependent transport processes. Na^+^-K^+^-2Cl^−^ symporter expression is unchanged at the 80%UL_50_ but increases 3-fold at the 80%LL_50_, corroborating a role in Cl^−^ uptake in dilute media.

*Halicarcinus plantus* shows a very different temporal expression profile for Na^+^/K^+^-ATPase α-subunit mRNA, unaltered over outset expression at the 80%UL_50_, suggesting a minor role in Cl^−^ secretion, but increasing 8-fold after 5 days at the 80%LL_50_, denoting a Na^+^/K^+^-ATPase-dependent Cl^−^ uptake mechanism. Unhappily, no data are available for Na^+^-K^+^-2Cl^−^ symporter mRNA expression in this species.

The time course of V(H^+^)-ATPase mRNA expression is unremarkable in both species with only minor differences. Expression is unaltered in *A. albatrossis*, but declines at the 80%UL_50_ in *H. planatus*, remaining unchanged at the 80%LL_50_, suggesting a negligible role for the V(H^+^)-ATPase at these levels of salinity challenge.

Together, these findings reveal that two sympatric, but distantly related Eubrachyura that occupy a coincident osmotic niche exhibit very similar systemic osmoregulatory characteristics such as their inability to regulate hemolymph osmolality and sodium concentration, yet strong chloride regulatory capability. While this might suggest convergent physiological adaptation molded by selection pressures in a common environment, the expressions of the transporter genes that typically underlie the mechanisms of ion uptake and secretion reveal considerable interspecific divergence. To illustrate, Na^+^/K^+^-ATPase expression is highly sensitive to low salinities, and V(H^+^)-ATPase expression to high salinities, in the hymenosomatid *Halicarcinus planatus*, but is modest in the belliid *Acanthocyclus albatrossis*. Apparently, the gene-based regulation of osmoregulatory processes has diversified between the two species, revealing disparity in the effects of likely similar selection pressures at different levels of structural organization, i. e., the genetic and the systemic. Further, both crabs appear to have limited regulation of their major hemolymph ions to Cl^−^ alone, suggesting an evolutionary trade-off between osmoregulatory energy expenditure and other temperature-dependent metabolic processes to which resources are apportioned.

## Acknowledgments

Crab collections in the Beagle Channel, Tierra del Fuego and the export of samples from Argentina to Brazil were authorized by the Secretaria de Ambiente, Ministerio de Producción y Ambiente. Provincia de Tierra del Fuego, Antártida e Islas del Atlántico Sur, Argentina (Permit # 63/2017) and a Material Transfer Agreement between CONICET and the Universidade de São Paulo. We are grateful to M. Torres, O. Florentín and C. Alonso for assistance with crab collections and fieldwork. We thank A. Giamportone and Dr Elaine Ribeiro for crab care and laboratory support at CADIC, and Susie Teixeira Keiko (DB, FFCLRP/USP) for technical assistance in Brazil. We are indebted to Dr. Ademilson Panunto Castelo (DB, FFCLRP/USP) for use of the CFX96 Real-Time PCR Detection System for qPCR analyses. JCM and ACM are especially grateful to the CADIC administration for providing excellent accommodation and thank the day staff for their warm hospitality.

## Data availability

All gene sequences generated in this study have been deposited with GenBank, National Center for Biotechnology Information, under the accession numbers provided. Other original data are available on reasonable request.

## Ethical approval

This investigation complies with all local, state, federal and international guidelines as regards the care and use of invertebrate animals in scientific research.

## References

Altschul, S. F., Gish, W., Miller, W., Myers, W. E. and Lipman, D. J. (1990). Basic local alignment search tool. J. Mol. Biol. 215, 403–410.

Augusto, A., Greene, L. J., Laure, H. J. and McNamara, J.C. (2007). Adaptive shifts in osmoregulatory strategy and the invasion of freshwater by brachyuran crabs: evidence from Dilocarcinus pagei (Trichodactylidae). J. Exp. Zool. A 307, 688–698.

Bedford, J. J. (1972). The composition of the blood of the grapsid crab, Helice crassa Dana. J. Exp. Mar. Biol. Ecol, 8, 113–119.

Benvides, H., Montoya, N. G., Carignan, M. and Luizón C. (2019). Environmental features and harmful algae in an area of bivalve shellfish production of the Beagle Channel, Argentina. Mar. Fish. Sci. 32, 71–101.

Bianchini, A., Lauer, M. M., Nery, L. E. M., Colares, E. P., Monserrat, J. M. and dos Santos Filho, E.A. (2008). Biochemical and physiological adaptations in the estuarine crab Neohelice granulata during salinity acclimation. Comp. Biochem. Physiol. A 151, 423–436.

Bozza, D. C., Freire, C. A. and Prodocimo, V. (2019). Osmo-ionic regulation and carbonic anhydrase, Na^+^/K^+^-ATPase and V-H^+^-ATPase activities in gills of the ancient freshwater crustacean Aegla schmitti (Anomura) exposed to high salinities. Comp. Biochem. Physiol. A 231, 201–208.

Bursey, C. R. (1982). Salinity tolerance and osmotic response in two species of spider crabs of the genus Libinia (Decapoda Brachyura, Majidae). Crustaceana. 194–200.

Cantelmo, A. C., Cantelmo, F. R. and Langsam, D. M. (1975). Osmoregulatory ability of the rock crab, Cancer irroratus, under osmotic stress. Comp. Biochem. Physiol. A 51, 537–542.

Capparelli, M. V., McNamara, J. C. and Grosell, M. (2017). Effects of waterborne copper delivered under two different exposure and salinity regimes on osmotic and ionic regulation in the mudflat fiddler crab, Minuca rapax (Ocypodidae, Brachyura). Ecotoxicol. Environ. Safety 143, 201–209.

Capparelli, M. V., Thurman, C. L., Choueri, P. G., de Souza Abessa, D.M., Fontes, M. K., Nobre, C. R. and McNamara, J. C. (2021). Survival strategies on a semi-arid island: submersion and desiccation tolerances of fiddler crabs from the Galapagos Archipelago. Mar. Biol. 168, 1–15.

Charmantier, G. and Charmantier-Daures, M. (1991). Ontogeny of osmoregulation and salinity tolerance in Cancer irroratus: elements of comparison with C. borealis (Crustacea, Decapoda). Biol. Bull. 180, 125–134.

Chen, X., Peng, Z., Hou, X., Wang, J. and Wang, C. (2019). The molecular basis of osmoregulation and physiological processes associated with salinity changes in the Chinese Mitten Crab Eriocheir sinensis. J. Shellfish Res. 38, 643–653.

Cieluch, U., Anger, K., Aujoulat, F., Buchholz, F., Charmantier-Daures, M. and Charmantie, G. (2004). Ontogeny of osmoregulatory structures and functions in the green crab Carcinus maenas (Crustacea, Decapoda). J. Exp. Biol. 207, 325–336.

Cornell, J. C. (1979). Salt and water balance in two marine spider crabs, Libinia emarginata and Pugettia producta. I. Urine production and magnesium regulation. Biol. Bull. 157, 221–233.

Curelovich, J., Lovrich, G. A. and Calcagno, J. A. (2009). New locality for Notochthamalus scabrosus (Crustacea, Cirripedia): Bahía Lapataia, Beagle Channel, Tierra del Fuego, Argentina. In: Anales del Instituto de la Patagonia. 37, 47–50.

D’Orazio, S. E. and Holliday, C. W. (1985). Gill Na, K-ATPase and osmoregulation in the sand fiddler crab, Uca pugilator. Physiol. Zool. 58, 364–373.

Dehnel, P. A. (1962). Aspects of osmoregulation in two species of intertidal crabs. Biol. Bull. 122, 208–227.

Diez, M. J., Cabreira, A. G., Madirolas, A., de Nascimento J. M., Scioscia, G., Schiavini, A. and Lovrich, G. A. (2018). Winter is cool: spatio?temporal patterns of the squat lobster Munida gregaria and the Fuegian sprat Sprattus fuegensis in a sub?Antarctic estuarine environment. Polar Biol. 41, 2591–260

Diez, M. J. and Lovrich, G. A. (2010). Reproductive biology of the crab Halicarcinus planatus (Brachyura, Hymenosomatidae) in sub-Antarctic waters. Polar. Biol. 33, 389–401.

Diez, M. J. and Lovrich, G. A. (2013). Moult cycle and growth of the crab Halicarcinus planatus (Brachyura, Hymenosomatidae) in the Beagle Channel, southern tip of South America. Helgol. Mar. Res. 67, 555–566.

Diez, M. J., Florentín, O. and Lovrich, G. A. (2011). Distribution and population structure of the crab Halicarcinus planatus (Brachyura, Hymenosomatidae) in the Beagle channel, Tierra del Fuego. Rev. Biol. Mar. Oceanogr. 46, 141–155.

Drach, P. and Tchernigovtzeff, C. (1967). Sur la méthode de détermination des stades d’intermue et son application générale aux crustacés. Vie et milieu, Serie A Biol. Mar. 29, 595–610.

Falconer, T. R., Marsden, I. D., Hill, J. V. and Glover, C. N. (2019). Does physiological tolerance to acute hypoxia and salinity change explain ecological niche in two intertidal crab species? Conserv. Physiol. 7, coz086.

Faleiros, R.O., Goldman, M.H.S., Furriel, R.P.M. and McNamara, J.C. (2010). Differential adjustment in gill Na^+^/K^+^- and V-ATPase activities and transporter mRNA expression during osmoregulatory acclimation in the cinnamon shrimp Macrobrachium amazonicum (Decapoda, Palaemonidae). J. Exp. Biol. 213, 3894–3905.

Faleiros, R. O., Furriel, R. P. and McNamara, J. C. (2017). Transcriptional, translational and systemic alterations during the time course of osmoregulatory acclimation in two palaemonid shrimps from distinct osmotic niches. Comp. Biochem. Physiol. A 212, 97–106.

Faleiros, R. O., Garçon, D. P., Lucena, M. N., McNamara, J. C. and Leone, F. A. (2018). Short-and long-term salinity challenge, osmoregulatory ability, and (Na^+^, K^+^)-ATPase kinetics and α-subunit mRNA expression in the gills of the thinstripe hermit crab Clibanarius symmetricus (Anomura, Diogenidae). Comp. Biochem. Physiol. A 225, 16–25.

Faria, S. C., Faleiros, R. O., Brayner, F. A., Alves, L. C., Bianchini, A., Romero, C., Buranelli R. C., Mantelatto, F. L. and McNamara, J. C. (2017). Macroevolution of thermal tolerance in intertidal crabs from Neotropical provinces: A phylogenetic comparative evaluation of critical limits. Ecol. Evol. 7, 3167–3176.

Faria, S. C., Provete, D. B., Thurman, C. L. and McNamara, J. C. (2017). Phylogenetic patterns and the adaptive evolution of osmoregulation in fiddler crabs (Brachyura, Uca). PloS one 12, e0171870.

Faria, S. C., Bianchini, A., Lauer, M. M., Zimbardi, A. L. R. L., Tapella, F., Romero, M. C. and McNamara, J. C. (2020). Living on the edge: Physiological and kinetic trade-offs shape thermal tolerance in intertidal crabs from tropical to sub-antarctic South America. Front. Physiol. 11, 312.

Finney, D.J., (1971). Probit Analysis, 3rd ed. Cambridge University Press, Cambridge.

Firmino, K. C., Faleiros, R. O., Masui, D. O., McNamara, J. C. and Furriel, R. F. (2011). Short- and long-term, salinity-induced modulation of V-ATPase activity in the posterior gills of the true freshwater crab, Dilocarcinus pagei (Brachyura, Trichodactylidae). Comp. Biochem. Physiol. B: Biochem. Mol. Biol. 160, 24–31.

Foster, C., Amado, E.M., Souza, M.M. and Freire, C.A. (2010). Do osmoregulators have lower capacity of muscle water regulation than osmoconformers? A study on decapod crustaceans. J. Exp. Zool. A 313, 80–94.

Frederich, M., Sartoris, F. and Pörtner, H O. (2001) Distribution patterns of decapod crustaceans in polar areas: a result of magnesium regulation? Polar Biol 24, 719–723.

Freire, C. A., Onken, H. and McNamara, J. C. (2008a). A structure-function analysis of ion transport in crustacean gills and excretory organs. Comp. Biochem. Physiol. 151, 272–304.

Freire, C. A., Amado, E. M., Souza, L. R., Veiga, M. P. T., Vitule, J. R. S., Souza, M. M., Prodocimo, V. (2008b). Muscle water control in crustaceans and fishes as a function of habitat, osmoregulatory capacity, and degree of euryhalinity. Comp. Biochem. Physiol. A 149, 435–446.

Freire, C. A., Souza-Bastos, L. R., Amado, E. M., Prodocimo, V. and Souza, M. M. (2013). Regulation of muscle hydration upon hypo-or hyper-osmotic shocks: differences related to invasion of the freshwater habitat by decapod crustaceans. J. Exp. Zool. A Ecol. Genet. Physiol. 319, 297–309.

Garçon, D. P., Masui, D. C., Mantelatto, F. L. M., Furriel, R. P. M., McNamara, J. C. and Leone, F. A. (2009). Hemolymph ionic regulation and adjustments in gill (Na^+^, K^+^)-ATPase activity during salinity acclimation in the swimming crab Callinectes ornatus (Decapoda, Brachyura). Comp. Biochem. Physiol. A 154, 44–55.

Garçon, D. P., Leone, F. A., Faleiros, R. O., Pinto, M. R., Moraes, C. M., Fabri, L. M., Antunes, C. D. and McNamara, J. C. (2021). Osmotic and ionic regulation, and kinetic characteristics of a posterior gill (Na^+^, K^+^)-ATPase from the blue crab Callinectes danae on acclimation to salinity challenge. Mar. Biol. 168, 1–19.

Genovese, G., Luchetti, C. G. and Luquet, C. M. (2004). Na^+^/K^+^-ATPase activity and gill ultrastructure in the hyper-hypo-regulating crab Chasmagnathus granulatus acclimated to dilute, normal, and concentrated seawater. Mar.Biol. 144, 111–118.

Gerencser, G.A. and Zhang, J.L. (2003) Chloride ATPase Pumps in Nature: do they exist? Biol. Rev. 78, 197–218.

Havird, J. C., Santos, S. R. and Henry, R.P. (2014). Osmoregulation in the Hawaiian anchialine shrimp Halocaridina rubra (Crustacea: Atyidae): expression of ion transporters, mitochondria-rich cell proliferation and hemolymph osmolality during salinity transfers. J. Exp. Biol. 217, 2309–2320.

Henry, R. P., Lucu, C., Onken, H. and Weihrauch, D. (2012). Multiple functions of the crustacean gill: osmotic/ionic regulation, acid-base balance, ammonia excretion, and bioaccumulation of toxic metals. Front. Physiol. 3, 1–33.

Hudson, D. M., Sexton, D. J., Wint, D., Capizzano, C. and Crivello, J. F. (2018). Physiological and behavioral response of the Asian shore crab, Hemigrapsus sanguineus, to salinity: implications for estuarine distribution and invasion. PeerJ 6, e5446.

Isla, F., Bujalevsky, G., Coronato, A. (1999) Procesos estuarinos em el Canal Beagle, Tierra del Fuego. Rev. Asoc. Geol. Argent. 54, 307–318

Jayasundara, N., Towle, D. W., Weihrauch, D. and Spanings-Pierrot, C. (2007). Gill-specific transcriptional regulation of Na^+^/K^+^-ATPase α-subunit in the euryhaline shore crab Pachygrapsus marmoratus: sequence variants and promoter structure. J. Exp. Biol. 210, 2070–2081.

Jillette, N., Cammack, L., Lowenstein, M. and Henry, R.P. (2011). Down-regulation of activity and expression of three transport-related proteins in the gills of the euryhaline green crab, Carcinus maenas, in response to high salinity acclimation. Comp. Biochem. Physiol. A 158, 189–193.

Leone, F. A., Garçon, D. P., Lucena, M. N., Faleiros, R. O., Azevedo, S. V., Pinto, M. R., McNamara, J. C. (2015). Gill-specific (Na^+^, K^+^)-ATPase activity and α-subunit mRNA expression during low-salinity acclimation of the ornate blue crab Callinectes ornatus (Decapoda, Brachyura). Comp. Biochem. Physiol. B 186, 59–67.

Leone, F. A., Lucena, M. N., Fabri, L. M., Garçon, D. P., Fontes, C. F., Faleiros, R. O., Moraes, C. M. and McNamara, J. C. (2020). Osmotic and ionic regulation, and modulation by protein kinases, FXYD2 peptide and ATP of gill (Na^+^, K^+^)-ATPase activity, in the swamp ghost crab Ucides cordatus (Brachyura, Ocypodidae). Comp. Biochem. Physiol. B 250, 110507.

Lin, H. C., Su, Y. C. and Su, S. H. (2002). A comparative study of osmoregulation in four fiddler crabs (Ocypodidae: Uca). Zool. Sci. 19, 643–650.

Livak, K. J. and Schmittgen, T. D. (2001). Analysis of relative gene expression data using real-time quantitative PCR and the 2-ΔΔCT method. Methods 25, 402–408.

López-Farrán, Z., Guillaumot, C., Vargas-Chacoff, L., Paschke, K., Dulière, V., Danis, B., Poulin, E., Saucède, T., Waters, J. and Gérard, K. (2021). Is the southern crab Halicarcinus planatus (Fabricius, 1775) the next invader of Antarctica? Glob. Change Biol. 27, 3487–3504.

Luquet, C. M., Weihrauch, D., Senek, M. and Towle, D.W. (2005). Induction of branchial ion transporter mRNA expression during acclimation to salinity change in the euryhaline crab Chasmagnathus granulatus. J. Exp. Biol. 208, 3627–3636.

Mantel, L. H. and Farmer, L. L. (1983). Osmotic and ionic regulation. In Mantel, L.H. (Ed.) The biology of Crustacea: 5. Internal anatomy and physiological regulation. The biology of Crustacea, pp. 53–161.

Mantovani, M. and McNamara, J. C. (2021). Contrasting strategies of osmotic and ionic regulation in freshwater crabs and shrimps: gene expression of gill ion transporters. J. Exp. Biol. 224, jeb233890.

Maraschi, A. C., Faria, S. C. and McNamara, J. C. (2021). Salt transport by the gill Na^+^-K^+^-2Cl-symporter in palaemonid shrimps: exploring physiological, molecular and evolutionary landscapes. Comp. Biochem. Physiol. A 257, 110968.

McNamara, J. C. and Faria, S. C. (2012). Evolution of osmoregulatory patterns and gill ion transport mechanisms in the decapod Crustacea: a review. J. Comp. Physiol. B 8, 997–1014.

McNamara, J. C. and Freire, C. A. (2022). Strategies of invertebrate osmoregulation: an evolutionary blueprint for transmuting into fresh water from the sea. BioRxiv preprint https://doi.org/10.1101/2022.01.16.476502.

McNamara, J. C. and Torres, A. H. (1999). Ultracytochemical location of Na^+^/K^+^-ATPase activity and effects of high salinity acclimation in gill and renal epithelia of the freshwater shrimp Macrobrachium olfersii (Crustacea, Decapoda). J. Exp. Biol. 284, 617–628.

McNamara, J. C., Freire, C. A., Torres, A. H. and Faria, S. C. (2015). The conquest of fresh water by the palaemonid shrimps: an evolutionary history scripted in the osmoregulatory epithelia of the gills and antennal glands. Biol. J. Linn. Soc. 114, 673–688.

Onken, H. and McNamara, J. C. (2002). Hyperosmoregulation in the red freshwater crab Dilocarcinus pagei (Brachyura, Trichodactylidae): structural and functional asymmetries of the posterior gills. J. Exp. Biol. 205, 167–175.

Péqueux, A. (1995). Osmotic regulation in crustaceans. J. Crust. Biol. 15, 1–60.

Pinto, M. R., Lucena, M. N., Faleiros, R. O., Almeida, E. A., McNamara, J.C. and Leone, F. A. (2016). Effects of ammonia stress in the Amazon river shrimp Macrobrachium amazonicum (Decapoda, Palaemonidae). Aquat. Toxicol. 170, 13–23.

Sanger, F., Nicklen, S. and Coulson, A. (1977). DNA sequencing with chain-terminating inhibitors. Proc. Natl. Acad. Sci. 74, 5463–5467.

Santos, F. H. and McNamara, J. C. (1996). Neuroendocrine modulation of osmoregulatory parameters in the freshwater shrimp Macrobrachium olfersii (Wiegmann) (Crustacea, Decapoda). J. Exp. Mar. Biol. Ecol. 206, 109–120.

Schales, O. and Schales, S.S. (1941). A simple and accurate method for the determination of chloride in biological fluids. J. Biol. Chem. 140, 879–883.

Siebers, D., Leweck, K., Markus, H. and Winkler, A. (1982). Sodium regulation in the shore crab Carcinus maenas as related to ambient salinity. Mar. Biol. 69, 37–43.

Smith, R. I. (1967). Osmotic regulation and adaptive reduction of water-permeability in a brackish-water crab Rhithropanopeus harrisi (Brachyura, Xanthidae). Biol. Bull. 133, 643–658.

Taylor, H. H. and Taylor, E. W. (1992). Gills and lungs: The exchange of gases and ions. In Microscopic Anatomy of Invertebrates, vol. 10 (ed. F. W. Harrison and A. G. Humes), pp. 203–293. New York: Wiley-Liss.

Thurman, C.L. (2002). Osmoregulation in six sympatric fiddler crabs (genus Uca) from the northwestern Gulf of Mexico. Mar. Ecol. 23, 269–284.

Thurman, C.L. (2003). Osmoregulation by six species of fiddler crabs (Uca) from the Mississippi delta area in the northern Gulf of Mexico. J. Exp. Mar. Biol. Ecol. 291, 233–253.

Thurman, C. L., Faria, S. C. and McNamara, J. C. (2017). Geographical variation in osmoregulatory abilities among populations of ten species of fiddler crabs from the Atlantic coast of Brazil: A macrophysiological analysis. J. Exp. Mar. Biol. Ecol. 497, 243–253.

Towle, D. W. and Weihrauch, D. (2001). Osmoregulation by gills of euryhaline crabs: molecular analysis of transporters. Am. Zool. 41, 770–780.

Tsai, J. R. and Lin, H. C. (2007). V-type H^+^-ATPase and Na^+^,K^+^-ATPase in the gills of 13 euryhaline crabs during salinity acclimation. J. Exp. Biol. 210, 620–627.

Wang, Z., Bai, Y., Zhang, D. and Tang, B. (2018). Adaptive evolution of osmoregulatory-related genes provides insight into salinity adaptation in Chinese mitten crab, Eriocheir sinensis. Genetica 146, 303–311.

Weihrauch, D., Ziegler, A., Siebers, D. and Towle, D. W. (2001). Molecular characterization of V-type H^+^-ATPase (B-subunit) in gills of euryhaline crabs and its physiological role in osmorregulatory ion uptake. J. Exp. Biol. 204, 25–37.

Weihrauch, D., McNamara, J.C., Towle, D.W. and Onken, H. (2004). Ion-motive ATPases and active, transbranchial NaCl uptake in the red freshwater crab, Dilocarcinus pagei (Decapoda, Trichodactylidae). J. Exp. Biol. 207, 4623–4631.

Weihrauch, D., Fehsenfeld, S. and Quijada-Rodriguez, A. (2017). Nitrogen excretion in aquatic crustaceans. In: Weihrauch, D., O’Donnell, M. (Eds.), Acid-Base Balance and Nitrogen Excretion in Invertebrates. Springer, Switzerland, Chapter 1, pp. 1–24

Whiteley, N. M., Suckling, C. C., Ciotti, B. J., Brown, J., McCarthy, I. D., Gimenez, L. and Hauton, C. (2018). Sensitivity to near-future CO2 conditions in marine crabs depends on their compensatory capacities for salinity change. Sci. Rep. 8, 1–13.

Zanders, I. P. and Rojas, W. E. (1996). Osmotic and ionic regulation in the fiddler crab Uca rapax acclimated to dilute and hypersaline seawater. Mar. Biol. 125, 315–320.

Zangrando, A. F., Ponce, J. F., Martinoli, M. P., Montes, A., Piana, E. and Vanella, F. (2016). Paleogeographic changes drove prehistoric fishing practices in the Cambaceres Bay (Tierra del Fuego, Argentina) during the middle and late Holocene. Environmental Archaeology: J. Human Paleoecol. 21, 182–192.

